# Integrated Molecular-Phenotypic Profiling Reveals Metabolic Control of Morphological Variation in Stembryos

**DOI:** 10.1101/2023.12.04.569921

**Authors:** Alba Villaronga Luque, Ryan Savill, Natalia López-Anguita, Adriano Bolondi, Sumit Garai, Seher Ipek Gassaloglu, Aayush Poddar, Aydan Bulut-Karslioglu, Jesse V Veenvliet

## Abstract

Mammalian stem-cell-based models of embryo development (stembryos) hold great promise in basic and applied research. However, considerable phenotypic variation despite identical culture conditions limits their potential. The biological processes underlying this seemingly stochastic variation are poorly understood. Here, we investigate the roots of this phenotypic variation by intersecting transcriptomic states and morphological history of individual stembryos across stages modeling post-implantation and early organogenesis. Through machine learning and integration of time-resolved single-cell RNA-sequencing with imaging-based quantitative phenotypic profiling, we identify early features predictive of the phenotypic end-state. Leveraging this predictive power revealed that early imbalance of oxidative phosphorylation and glycolysis results in aberrant morphology and a neural lineage bias that can be corrected by metabolic interventions. Collectively, our work establishes divergent metabolic states as drivers of phenotypic variation, and offers a broadly applicable framework to chart and predict phenotypic variation in organoid systems. The strategy can be leveraged to identify and control underlying biological processes, ultimately increasing the reproducibility of in vitro systems.

**Highlights:**

- Time-resolved single-cell RNA-sequencing and imaging-based quantitative charting of hundreds of individual stembryos generates molecular and phenotypic fingerprints
- Machine learning and integration of molecular and phenotypic fingerprints identifies features and biological processes predictive of phenotypic end-state
- Early imbalance of oxidative phosphorylation and glycolysis results in aberrant morphology and cellular composition
- Metabolic interventions tune stembryo end-state and can correct derailment of differentiation outcomes

## INTRODUCTION

As mammalian embryos develop *in utero*, direct observation and manipulation of developmental processes is difficult. This obstacle can be overcome by coaxing human and mouse pluripotent stem cells (PSCs) to self-organize into three-dimensional structures, called stembryos, reflecting many molecular, cellular and morphological characteristics of the embryo (stembryo: a portmanteau of stem cells and embryos).^1,2^ Because stembryos are easy to access, track, manipulate and scale, more quantitative and detailed mechanistic insights can be obtained in statistically relevant sample sizes. Hence, they offer huge potential in both fundamental (e.g. genetic screens, lineage tracing) and applied (e.g. drug screenings, embryo toxicology) research.^3^ However, a major hurdle for the widespread implementation of stembryos is considerable phenotypic variation that is largely unexplained.^1,2,4^ More broadly, inter-structure variation, presumably arising from genetic heterogeneity, cellular interactions, and the micro-environment, poses a significant challenge in organoid research.^5,6^

Consequently, unraveling the biological processes that underlie phenotypic variation is crucial to maximize the potential applications of organoids and stembryos. This remains a formidable task, especially since conventional assays to infer biological processes in an unbiased manner (e.g. single-cell RNA-sequencing (scRNA-seq)) necessitate the destruction of samples, making it impossible to simultaneously capture both the *current* gene expression state and the *future* phenotypic state within the *same* sample. Hence, a framework that enables integrated, longitudinal analysis of gene expression and phenotypic dynamics (phenodynamics) is required to obtain insights into the roots of divergent differentiation outcomes *in vitro*.

Here, we combined high-throughput longitudinal imaging and scRNA-seq of individual structures, quantitative image analysis, and machine learning, to map the expression and phenotypic landscapes of individual gastruloids and trunk-like structures.^7–11^ By integrating time-resolved expression profiles with phenodynamics, we successfully identified early predictors of stembryo end-states. Through sampling across these predictive phenotypic spaces, we uncovered biological processes strongly associated with phenotypic variation. Specifically, balanced glycolysis and oxidative phosphorylation in neuromesodermal progenitors (NMPs^12–14)^ underlies the harmonious development of somitic and neural tissues and modulation of metabolic activity can effectively tune stembryo differentiation outcomes. Collectively, our findings establish divergent metabolic states as drivers of phenotypic variation in stembryos.

## RESULTS

### Divergent differentiation outcomes in stembryos modeling the embryonic trunk

Gastruloids are aggregates of mouse embryonic stem cells (mESCs) that break symmetry, elongate and self-organize the major body axes upon a 24h-pulse with the WNT agonist CHIR99021 (hereafter CHIR) (**Figure 1A**).^7,11,8,9^ When supplemented with the extracellular matrix compound Matrigel, gastruloids develop into trunk-like-structures (TLSs) resembling the core part of the embryonic trunk (**Figure 1A**).^10,15^ While nearly half of the TLSs exhibit coordinated formation of somites and a neural tube, why >50% of the structures fail to develop properly diversified and organized tissues is unclear (**Figure 1A,B**).^10,15^ We therefore investigated the complete TLS morphospace in more detail upon initial classification of TLSs based on bright-field imaging (“successful” and “unsuccessful” – see Methods for criteria^10^). To ease visualization and quantification, we generated structures from mESCs carrying a T::H2B-mCherry (hereafter T^mCH^) and Sox2::H2B-Venus (hereafter Sox2^VE^) reporter, labeling cells actively expressing T (also known as Brachyury; expressed in NMPs and nascent mesoderm) or Sox2 (expressed in NMPs and their neural descendants) as well as their progeny.^10,16^ Overall architecture was assessed by DAPI (nuclei) and Phalloidin (F-Actin) staining. While successful TLSs form a central neural tube-like structure flanked by somites, unsuccessful TLSs showed disorganized neural tissue, or, in few cases, internalization of mesodermal tissue enveloped by neural tubes (**Figure 1C-E and S1A,B**). Accordingly, in unsuccessful structures the T^mCH+^ volume fraction, but not intensity, is reduced (**Figure 1C,D and S1C,D**).

**Figure 1.**
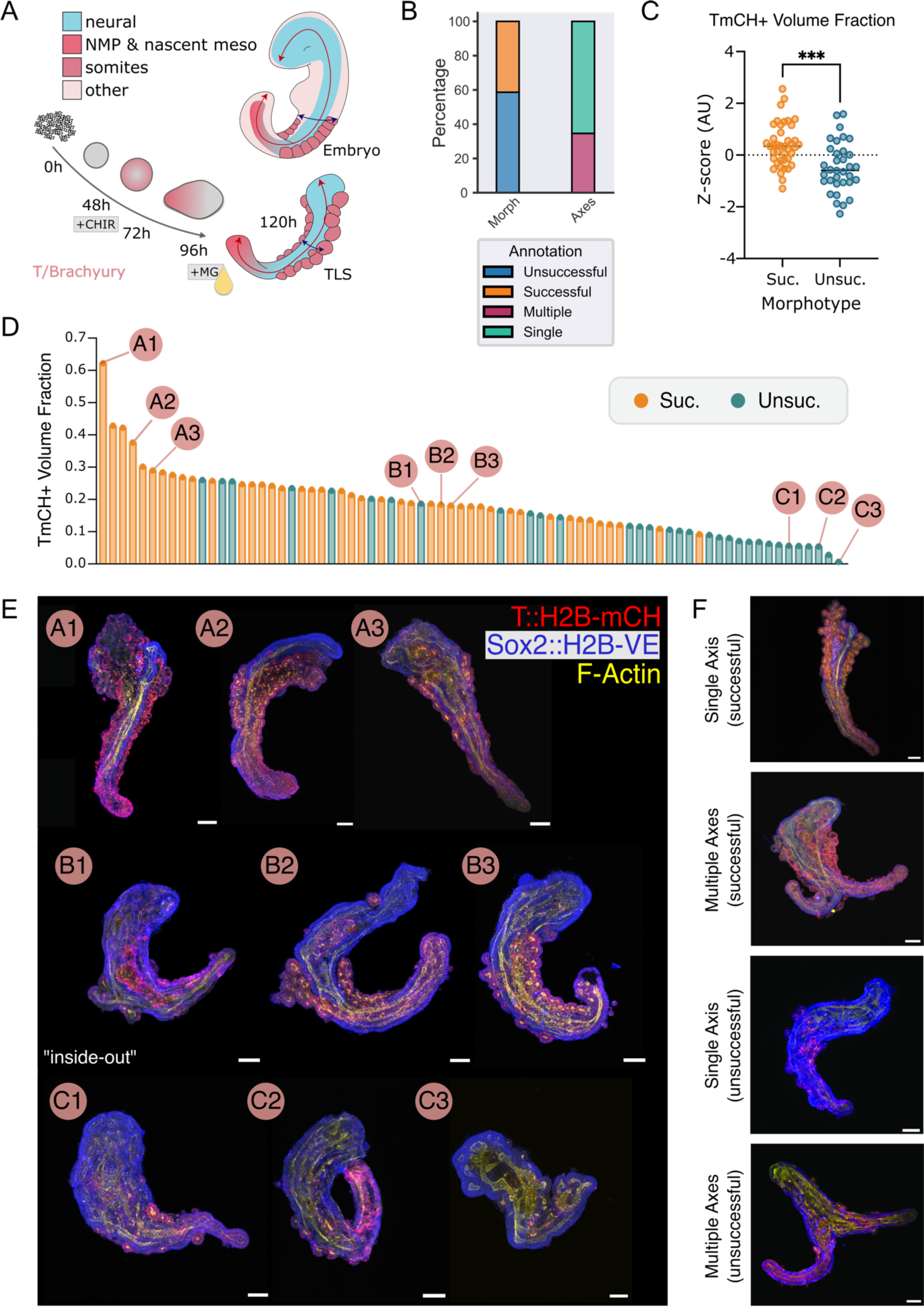
Divergent differentiation outcomes in trunk-like-structures. (A) Schematic overview of TLS formation (see Methods) and comparison with the embryo. Note that TLSs model the core part of the embryonic trunk (posterior neural tube + somites). (B) Quantifications of indicated morphological features in TLSs (scoring criteria in^10^). (C) T^mCH+^ Volume Fraction quantification of successful (n=41) and unsuccessful (n=34) unipolar TLSs. ***, p < 0.001 (D) Histogram showing distribution of T^mCH+^ Volume Fraction in TLSs. (E) Representative 3D maximum intensity projections (3D-MIPs) of unipolar TLSs with high (A1-A3), middle (B1-B3) and low (C1-C3) T^mCH+^ volume fraction. Scale bar, 100µm. (F) Representative 3D-MIPs of successful or unsuccessful unipolar and multipolar TLSs. Scale bar, 100µm.

A second frequently observed aberrant phenotype (compared to the embryo) was the formation of multi-axes structures with a total volume comparable to those with a single axis (**Figure 1B,F and S1E**).^10^ Notably, in multipolar structures without morphological somite formation the T^mCH+^ volume fraction was decreased less than in unipolar structures (**Figure S1F**).

### Quantitative charting of phenodynamics

We hypothesized that subtle phenotypic variation at earlier stages could precede divergent end-state differentiation outcomes. To test this, we quantitatively charted the 48-96h stembryo phenodynamics of a total of 768 stembryos and performed wide-field imaging at 24 hour-intervals (**Figure 2A**). Using deep-learning-based segmentation and a combination of feature extraction approaches^17–21^, we extracted: i) structure-level morphometric features from the bright-field (BF) data (hereafter referred to as *[feature]*-BF); ii) morphometric and distribution features from the T^mCH+^ domain (hereafter referred to as *[feature]*-T^mCH^); iii) intensity features measured as a function of the T^mCH+^ domain or whole structure (hereafter referred to as *[feature]*-T^mCH(mCH)^ and *[feature]*-T^mCH(BF)^ respectively) (**Figure 2A**) (**Supplementary Note 1**).

**Figure 2.**
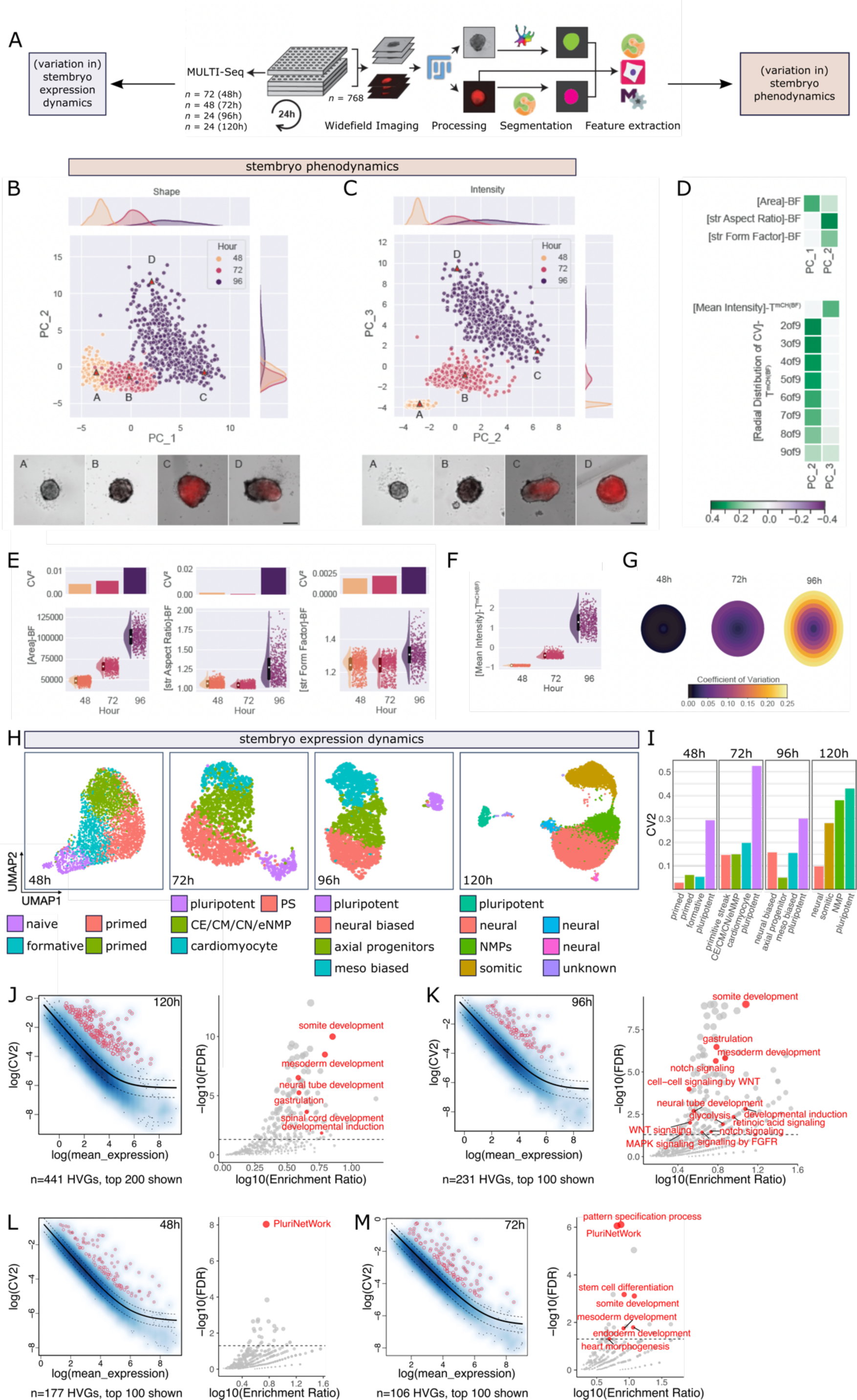
Time-resolved charting of the stembryo phenotypic and transcriptional landscape. (A) Processing pipeline for wide-field data. From 48h onwards, every 24 all structures are imaged and selected structures are processed for single-cell RNA-sequencing (scRNA-seq). After pre-processing and segmentation, features are extracted using scikit-image, CellProfiler and MOrgAna. (B,C) Scatterplots of principal components determined using sparse PCA (sPCA) with BF (B) and T^mCH^ (C) features of all timepoints. Timepoints are color-coded to visualize the phenodynamics and variation in the dataset. Top and left of the scatter-plot density plots of the x- and y-axis are shown. Example structures are annotated and shown in the bottom panels. (D) Feature loadings of principal components shown in B and C. Top: BF feature sPCA loadings, bottom: T^mCH^ feature sPCA loadings. (E,F) Plots of individual features with high loadings for BF feature sPCA (E) and [feature]-T^mCH^ sPCA (F). Bar plot (top) shows *CV*^2^ of features at each timepoint; raincloud plot (bottom) shows distribution of indicated features per time-point. Errorbar in raincloud plot indicates standard deviation and white center shows mean value. (G) *[Radial Distribution of CV]*-T^mCH(BF)^ visualization. Structures are approximated as ellipses with average major and minor axes at the respective timepoints. Concentric ellipses represent the radial segments in which CV values are colour-coded. (H) Uniform Manifold Approximation and Projection (UMAP) visualisations of scRNA-seq expression profiles of pooled individual stembryos at indicated time-points, coloured by cluster. (I) Bar plots showing *CV*^2^ for cell composition, computed for each cluster at indicated time-points. Colour-coding of clusters matches H. (J-M, left panels) Scatter plot showing mean and variance of individual genes (black dots) at indicated time-points. A generalized linear model was fitted to the data to capture the relationship between mean and variance. Highly variable genes (subset shown in red) were identified based on this model. (J-M, right panels) Results of Over-Representation Analysis (ORA). Selected terms of interest are highlighted in red. Sample sizes: B,E,F: 48h, n=766; 72h, n=662; 96h, n=546, C,G: 48h, n=545; 72h, n=546; 96h, n=546. In B,C,E,F only values > 0.005^th^ quantile and < 0.995^th^ quantile are plotted.

We then employed unsupervised machine learning in the form of Sparse Principal Component Analysis (SPCA) (**Figure 2B,C**). With *[feature]*-BF as input the first principal component (PC1) separated stembryos based on culture time; loadings were dominated by area and minor axis length, and features strongly correlated with both (**Figure 2B,D and S2A,C,D**). PC2 comprised *[AR]*-BF (*A*spect *R*atio), *[eccentricity]*-BF and first Hu-moment (*[hu1]*-BF - a scale, rotation and translation invariant description of shape) (**S2D,E**). Consequently, 96h structures were separated from other timepoints, reflecting the onset of axial elongation (**Figure 2B,D and S2E**). A second major contributor, [*form factor*]-BF, reflects the emergence of complex shapes around 96h (**Figure 2D**).

To quantify phenotypic variation, we calculated the squared coefficient of variation (*CV*^2^) for the dominant contributors over time, which showed that phenotypic variation overall increases as stembryos develop (**Figure 2E**). Decreased *CV*^2^ for a subset of features (*[AR]*-BF, *[hu1]*-BF) from 48-72h suggests that some variation emerging during aggregation is constrained by the CHIR pulse (**Figure 2E and S2E**).

Similarly, SPCA with *[feature]*-T^mCH^ as input sorted stembryos along a temporal trajectory, especially when considering PC2 and PC3 (**Figure 2C**; PC1 discussed in **Supplementary Note 2**). Evaluation of PC3 loadings showed a high contribution of *[mean_intensity]*-T^mC(BF)^, which steadily increased over time (**Figure 2D,F**). Other PC3 loadings correlated significantly with *[mean_intensity]*-T^mCH(BF)^ (**Figure S2F,G**). PC2 loadings were dominated by the coefficient of variation (*CV*) calculated across the radial slices of the mask (*[radial_distribution_CV]*-T^mCH(BF)^), a proxy for the homogeneity of intensity in the core versus periphery (**Figure 2G; Supplementary Note 2**). Hence, higher *CV* values in the periphery indicate T^mCH^ polarization over time. Indeed, PC4 separated 96h structures from earlier timepoints, with polarization along the major and minor axes as predominant loadings (**Figure S2F,I**).

In conclusion, phenodynamic analysis could capture the previously described growth, elongation, and establishment of polarized Brachyury expression.^7–11,22^ In addition, it revealed that phenotypic variation increases over time.

### Time-resolved single-cell RNA-sequencing across the phenotypic landscape

In parallel with phenodynamic analysis, we assessed heterogeneity in gene expression dynamics and cell state composition. We used MULTI-Seq^23^ to profile individual stembryos by scRNA-seq at 48h (n=72), 72h (n=48), 96h (n=24) and 120h (n=24) (**Figure 2A and S3A-C**). After stringent quality control (see Methods), we recovered high-quality transcriptomes of 21,026 cells (48h: 3,861 cells; 72h: 1,742 cells, 96h: 2,056 cells; 120h: 13,367 cells).

First, we aggregated all structures on a per time-point basis to obtain a time-resolved compendium of stembryo development. Inspection of cluster marker genes, intersection with data capturing naïve-to-primed pluripotency transitions *in vitro*, and pseudotime analysis showed that 48h structures harbor cells in naïve, formative and primed pluripotency states, whereas lineage-specific factors are not expressed (**Figure 2H and S3D-H**).^24,25^ 24 hours later, most cells had acquired signatures of the primitive streak, caudal epiblast, caudal mesoderm, caudal neuroectoderm, nascent mesoderm and cardiomyocytes (annotations based on embryo data^26^) (**Figure 2H and S3D’**). Pseudotime inference showed a trajectory from a naïve pluripotent state, via an epiblast and primitive streak signature, to an axial progenitor (caudal epiblast, mesoderm and neuroectoderm) state (**Figure S3J,K**).

Higher-level separation at 96h demonstrated a small partition with a pluripotency signature, and a larger one comprising three clusters (**Figure S3L**), organized into a continuum of states, with axial progenitors ((early) NMPs) flanked by mesodermal or neural biased cells (**Figure 2H and Figure S3D’’,M,N**). A similar organization was observed at 120h, with NMPs adjoined by somitic and neural cells (**Figure 2H and S3D’’’,O,P**). Notably, fine-grained clustering of the pluripotent cluster revealed two distinct populations, characterized by high expression of naïve and formative pluripotency genes, and primordial germ cell (PGC) marker genes respectively (PGC-like cells (PGCLCs)) (**Figure S3Q-S**). These two states were detected as early as 48h (**Figure S3S,T**). Hence, the recently identified ectopic pluripotent cells (EPCs) in gastruloids are also present in TLSs.^27^

### Profiling of inter-stembryo transcriptional variation

We leveraged the single-cell, single-structure resolution of our data-set to assess inter-structure variation in gene expression and cellular composition. The latter varied considerably between stembryos at all timepoints, with inter-structure variation increasing over time (**Figure 2I and Figure S4A-D**). At 120h unsuccessful and successful structures significantly differed in the fraction of somitic and neural cells, providing further support for our manual morphology-based classification (**Figure 1 and S4E**).

We then computed highly variable genes (HVGs) across individual structures (see Methods) (**Figure 2J-M and S4F-I**). As expected, at the end-state (120h) HVGs were involved in somite, mesoderm, neural tube and spinal cord development and their developmental induction (**Figure 2J and S4I,K**). Variation in the expression of genes involved in these processes was already prominent at 96h, when marker genes of NMPs (e.g. *Epha5*, *Cdx2/4*) as well as their neural (e.g. *Irx1/2/5*, *Sox1*, *Dbx1*, *Rfx4*) and mesodermal (e.g. *Uncx*, *Foxc2*, *Tcf15*, *Pcdh8*) progeny displayed significantly more variation than randomly selected or all other expressed genes (**Figure 2K and S4H,J**). Concomitantly, expression of gene modules for signaling pathways involved in axial patterning, NMP maintenance and/or decision making (FGF, WNT, Retinoid Acid, Notch) showed high inter-structure variation (**Figure 2K**). While at earlier time-points (48h and 72h), HVGs were dominated by pluripotency genes (**Figure 2L,M and S4F,G**), inter-structure divergence in mesoderm and somite development signatures became evident at 72h (**Figure 2M**).

The variation in cell state composition suggested extensive inter-stembryo heterochronicity. Indeed, pseudotime distributions were highly variable between individual structures (**Figure S4L-O**). To identify genes associated with heterogeneous developmental time, we correlated pseudo-bulked expression with pseudotime values. At 48h, “younger” and “older” structures showed enriched expression of naïve (e.g. *Klf2, Fbxo15)* whereas “older” and formative (e.g. *Sox4*, *Dnmt3b*) pluripotency markers respectively (**Figure S4P**), confirming the validity of our approach. At 72h, less advanced developmental time was associated with high expression of epiblast genes (e.g. *Utf1*, *Lefty1/2* and *Tdgf1*), while developmentally more progressed structures showed increased expression of primitive streak genes (e.g. *Gbx2*, *Hoxa1*) (**Figure S4Q**). Younger 96h gastruloids were characterized by high expression of primitive streak and axial progenitor markers (including *T*), whereas older gastruloids displayed higher expression of neural genes (**Figure S4R**). Finally, at 120h, more advanced developmental progression was associated with an increased neural expression signature at the expense of NMP marker genes (**Figure S4S**).

In sum, these data show that molecular and phenotypic variation emerge concomitantly. Next, we set out to integrate both modalities.

### Phenotypic fingerprints enable early prediction of morphological end-state

The destructive nature of scRNA-seq made it impossible to simultaneously capture both the *current* molecular and *future* phenotypic state within the *same* sample, prohibiting their integration. To overcome this, we aimed to identify early phenotypic features predictive of the stembryo end-state, in order to correlate early expression profiles to (predicted) morphological outcome.

We used two different approaches: i) integration of end-state expression with phenodynamics data (next section); ii) Partial Least Squares Regression (PLSR), a method designed to determine a latent space in which the explanatory and target variables show high covariance.^28,29^ This space is determined by a set of latent variables combinations of features similar to principal components, which can be analyzed in terms of their abilities to separate phenotypic (end-)states and the contributing features.^29,30^

When using differentiation outcomes as the target vector, the first latent variable (Dev-Dim1) clearly separated successful and unsuccessful TLSs (**Figure 3A**). Examining its loadings revealed significant contributions of 72h and 96h T^mCH^ levels, along with the T^mCH^ radial distribution features (**Figure 3C**) and highly correlated features (**Table S1**). At both 72h and 96h high(er) T^mCH^ levels were strongly associated with successful TLS formation, in particular in unipolar structures (**Figure 3C,E-G**). In terms of distribution, higher T^mCH^ levels at the core were associated with better differentiation outcome, as demonstrated by computing “virtual” average gastruloids (**Figure 3C,D**). These simplified representations also showed that structures with an increased T^mCH+^ area fraction at 96h (96h-*[fraction]*-T^mCH^) exhibited better differentiation outcomes in unipolar structures (**Figure 3D,H**).

**Figure 3.**
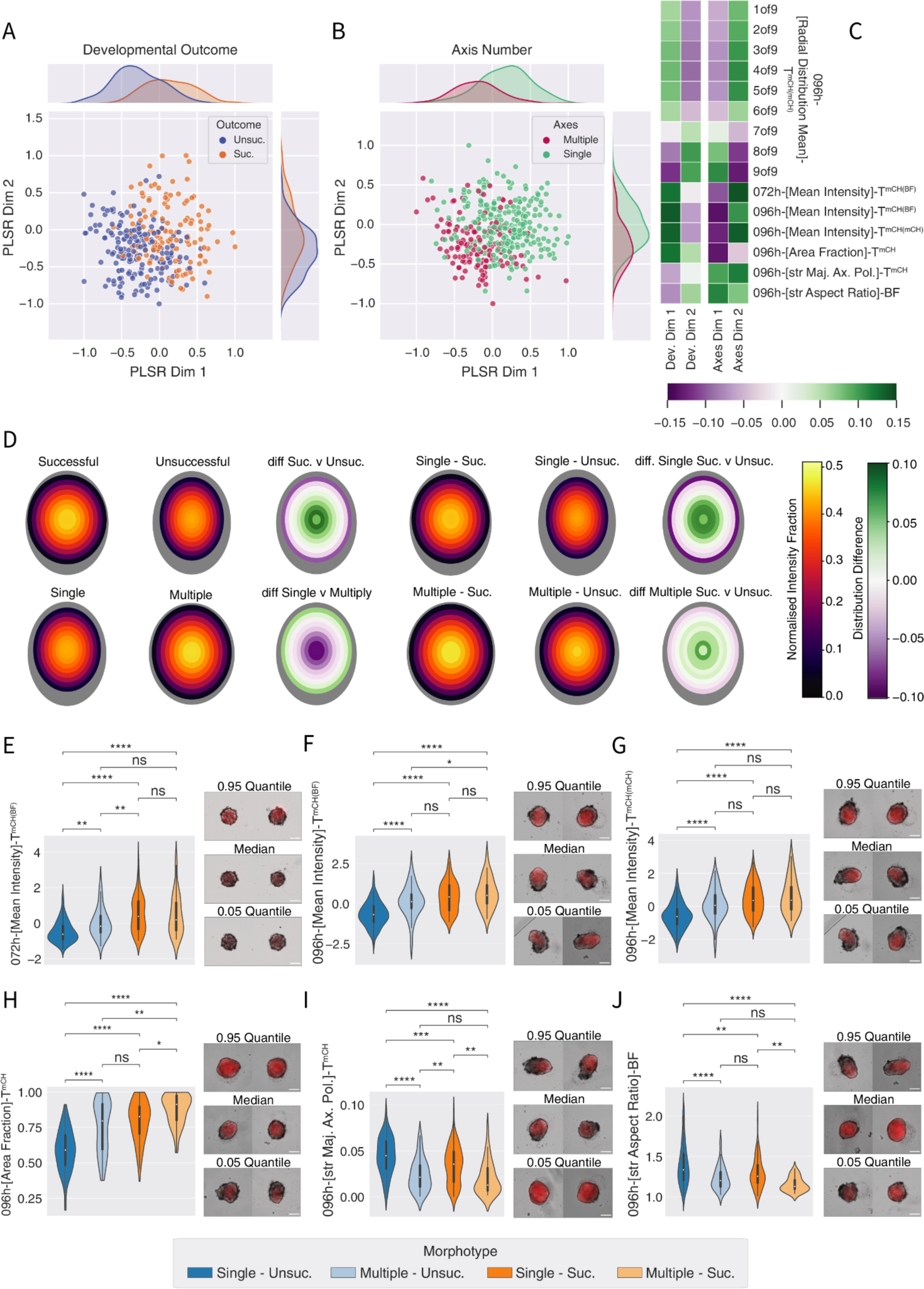
Identification of phenotypic features associated with stembryo differentiation outcome. (A,B) Scatterplots of PLSR latent variables. To the top and right of the plots the density plots of the x and y axis values, respectively are shown. Suc, Successful; Unsuc, unsuccessful. (C) Loading weights of features shown in A and B for both developmental outcomes (left: Dev. Dim 1, Dev. Dim 2) and axis number latent variables (right: Axes Dim 1, Axes Dim 2). (D) 96h-[*Radial Distribution of CV*]-T^mCH(mCH)^ visualization. Outer ellipses: approximation of structure shape with average major and minor axes; Inner ellipses: approximation of T^mCH^ domain shape with average major and minor axes. Inner concentric ellipses represent the radial segments in which the normalized intensity fraction is visualized by color. Distribution difference visualization - Inner concentric ellipses represent the radial segments in which the difference between morphotypes is visualized. (E-J) Left panels: violin plots of indicated features. * p < 0.05, ** p < 0.01, *** p < 0.001, **** p < 0.0001. Right panels: example structures with values closest to the 0.95 quantile (top), median value (middle) and 0.05 quantile (bottom). Images show composites of BF (greyscale) and T^mCH^ (red). Scale bars, 200µm. Sample sizes: Sample sizes A: Unsuccessful, n=183, Successful, n=128. Sample sizes B: Multiple, n=136; Single, n=256. D-J: Unsuccessful, n=183, Successful, n=129, Multiple, n=136; Single, n=257; Multiple-Unsuccessful, n=55; Multiple-Successful, n=34; Single-Unsuccessful, n=128; Single-Successful, n=95.

When investigating the first latent variable for axes number as the target vector (Axes-Dim1), many features identified for differentiation outcome reappeared as major contributors (**Figure 3B,C and Table S1**). Specifically, for 120h single-axis structures T^mCH^ intensity at 72h and 96h was overall lower, higher in the periphery versus the core of the T^mCH^ domain, and more polarized (**Figure 3C-I**). Compared to the differentiation outcome PLSR results, there was a higher contribution of 96h BF shape descriptors to Axes-Dim1; more elongated structures were more likely to become unipolar (**Figure 3I,J**).

We next tested the feasibility to predict phenotypic (end-)state at earlier time-points. Reduced feature sets comprising only 48h and 72h features showed convincing separation of phenotypic end-states (**Figure S5A,B**). For axes number, predictive features shifted from shape to T^mCH^ intensity and distribution features (**Figure S5C,D and Table S1**). Similar to the analysis using the complete feature space, higher 72h-*[mean_intensity]*-T^mCH(BF)^ was associated with better differentiation outcome (**Figure 3E and Table S1**), and T^mCH^ signal distribution with both axes number and differentiation outcome (**Figure S5C,D and Table S1**).

Overall, the PLSR analysis identified early phenotypic features that harbor predictive power for stembryo phenotypic end-state. In particular, successful TLS formation is strongly associated with: i) higher activation of the T^mCH^ reporter at 72h and 96h; ii) a higher fraction of the structure expressing T^mCH^ at 96h; iii) a higher reporter signal in the core compared to the periphery at 96h. For single versus double axes, both overall shape (e.g. AR, minor axis length), as well as T^mCH^ intensity and distribution harbor predictive power.

### Integrated molecular-phenotypic analysis identifies features predictive of differentiation outcome

Although imaging-based analysis alone identified predictive features, our analysis of the phenotypic landscape showed that for differentiation outcome the data followed a continuum rather than discrete distribution (**Figure 1D,E**). We therefore leveraged that our experimental design comprised parallel recording of phenotypic and transcriptional states, including ground truth outcomes.

Specifically, we aimed to identify early phenotypic features predictive of end-state somitic vs neural ratio. We computed somitic and neural module scores for individual 120h structures and used these scores as “bait” in a correlation analysis with the phenotypic features (**Figure 4A and S5E**). This integration of both high-dimensional spaces (i.e. 48h/72h/96h phenotypic space and 120h transcriptional space) revealed high absolute correlations of somitic and neural module scores with T^mCH^ intensity and distribution, but not BF, features (**Figure 4B,C and S5F**). Specifically, we observed a high positive correlation between the 120h somitic module score and both 96h-*[fraction]*-T^mCH^ and 96h-*[mean_intensity]*-T^mCH(mCH)^. Vice versa, the neural module score was strongly negatively correlated with these features (**Figure 4B,C and Figure S5G**). These associations were also apparent upon quartile binning of structures based on *[fraction]*-T^mCH^ or *[mean_intensity]*-T^mCH(mCH)^; with higher values, the percentage of NMP and somitic cells increased, and cells occupied distinct regions of the 120h compendium’s expression space (**Figure 4D,E and Figure S5H**). A small set of 72h T^mCH^ features was strongly associated with future differentiation outcome; in particular 72h-*[mean_intensity]*-T^mCH^ correlated with an increased somitic vs neural fraction at 120h, as well as a relative increase of NMPs (**Figure 4B,C,F**).

**Figure 4.**
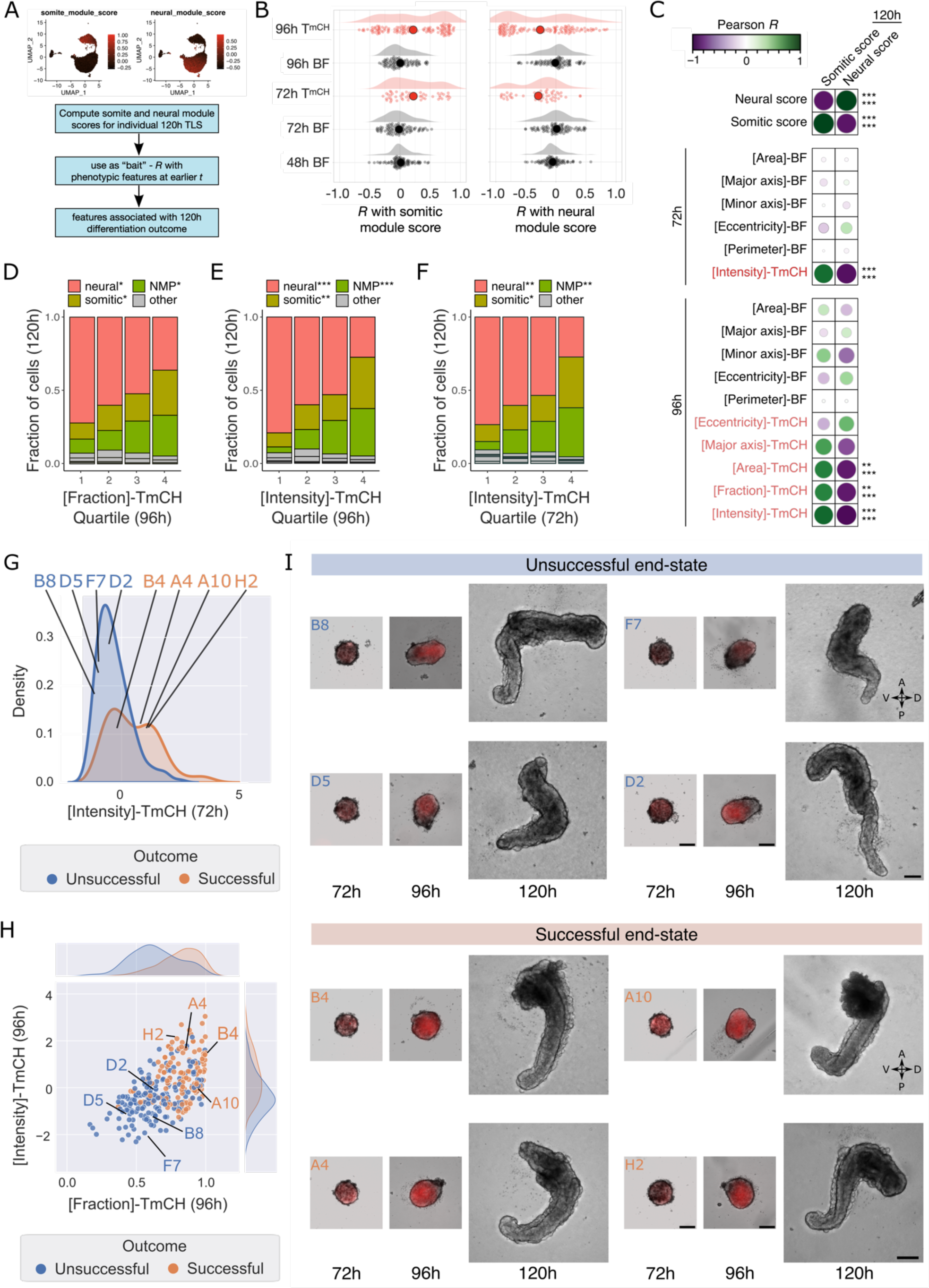
Integration of molecular and phenotypic fingerprints identifies predictive features. (A) Schematic of computational strategy to identify predictive features through integration of imaging and scRNA-seq data. (B) Raincloud plot showing the distribution of computed correlations of grouped feature sets (on y-axis) with somitic (left) or neural (right) module scores. BF, brightfield. (C) Dot plots showing correlations for selected features. ** padj < 0.01, *** padj < 0.001 (Holm-Bonferroni correction for multiple testing). Top: padj for correlation with somitic score, bottom: padj for correlation with neural score. (D-F) Cumulative bar plot showing the fraction of cells at 120h in each color-coded annotated state for (D) 96h [Fraction]-T^mCH^ quartiles; (E) 96h [Intensity]-T^mCH^ quartiles; (F) 72h [Intensity]-T^mCH^ quartiles. * p<0.05, ** p<0.01, *** p<0.001 (ANOVA with Benjamini-Hochberg FDR correction). NMP, neuro-mesodermal progenitor. (G) Density plot showing distribution of [Intensity]-T^mCH^ at 72h for structures with successful (orange) and unsuccessful (blue) outcomes at 120h. (H) Simplified feature space showing distribution of [Intensity]-T^mCH^ and [Fraction]-T^mCH^ at 96h for structures with successful (orange) and unsuccessful (blue) outcomes at 120h. Example structures (annotated in density plot) are shown in the right panel. Scale bars, 200 µm. (I) Examples of structures (annotated in G and H) predicted to give rise to unsuccessful (top) and successful (bottom) unipolar TLSs, including ground truth end-state. Scale bars, 200 µm. A, anterior; P, posterior; D, dorsal; V, ventral.

To assess the predictive power, we compared machine learning classifiers of varying complexity (Linear Discriminant Analysis (LDA)^31^, State Vector Classifier (SVC)^32^ and eXtreme Gradient Boosting (XGBoost)^33^) in their ability to predict axis number and differentiation outcome based on all features or predictive features only. SVC and XGBoost displayed similar accuracies, whereas LDA scored lower (**Figure S5I,J**). Whereas feature selection only marginally impacted SVC and XGBoost classification scores, LDA benefited, possibly due to the reduced complexity of its decision surface (**Figure S5I,J**).

Altogether, our integrated molecular-phenotypic analysis demonstrated that 96h-*[fraction]*-T^mCH^ and 72h- and 96h-*[mean_intensity]*-T^mCH^ are strongly associated with differentiation outcomes at 120h. This, together with the PLSR analysis (**Figure 3**), suggests that these features are dominant predictors of the structure’s ability to balance somite and neural tube formation. Indeed, projecting the 120h phenotypic outcome on a simplified 72h or 96h feature space well-separated 120h successful and unsuccessful structures, showing that these features can predict the stembryo end-state (**Figure 4G-I**).

### Identification of divergent glycolytic activity as a major putative driver of variation

Next, we utilized the newly obtained predictive power to identify biological processes involved in divergent differentiation outcomes. The even distribution of sequenced 96h structures across the predictive state space allowed us to match these with phenotypically similar structures for which the ground truth end-state was captured (**Figure 5A,B**). As discussed above, on average structures at this stage comprised 4 major cell states: pluripotent cells, and axial progenitors (early NMPs) flanked by mesodermal or neural biased cells (**Figure 2H**). This lineage bias was reflected by predictive phenotypic features (**Figure S6A,B**).

**Figure 5.**
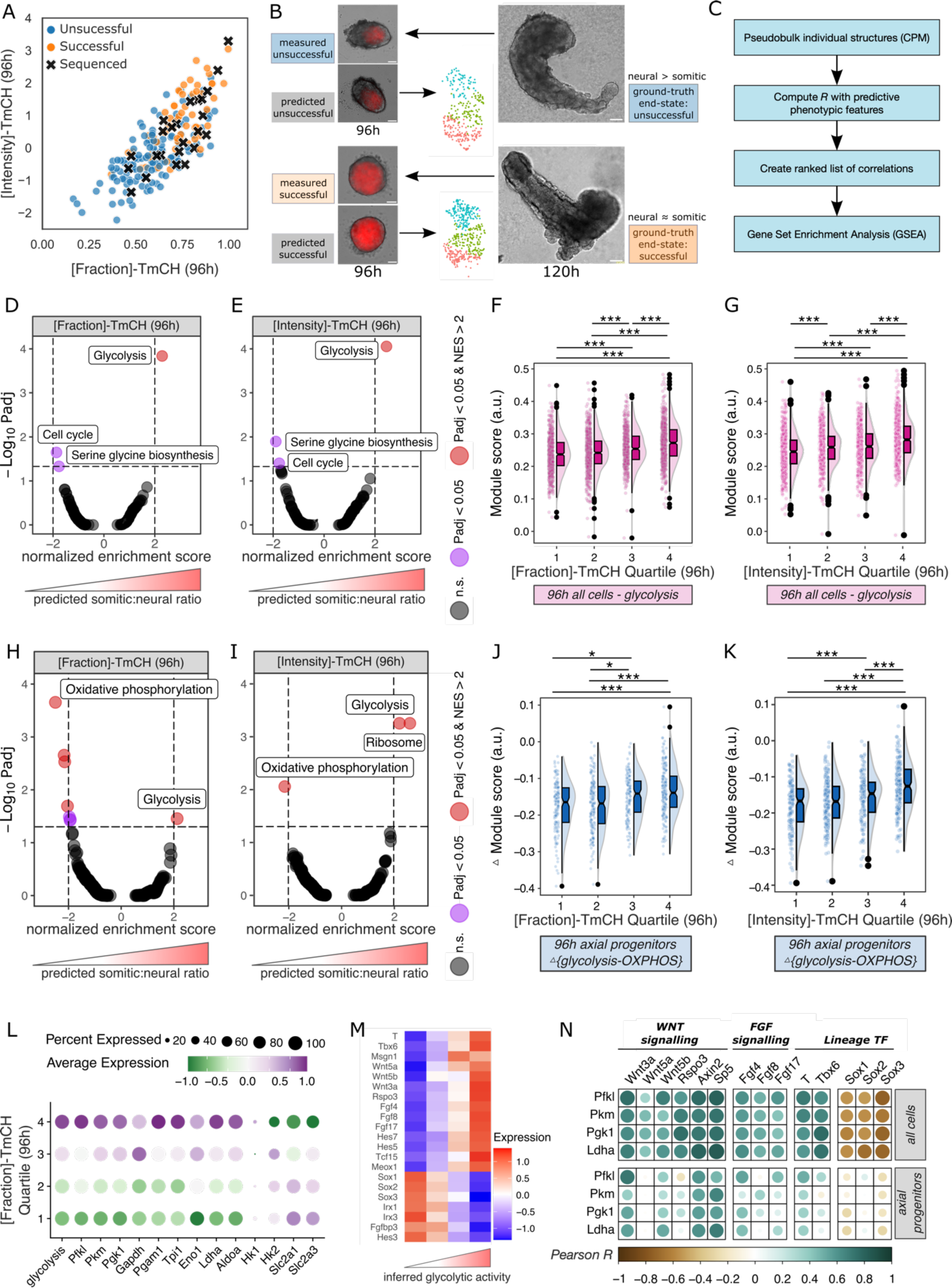
Early activity of glycolysis and OxPhos predicts divergent differentiation outcome. (A) Scatter plot showing distribution of 96h sequenced structures (black crosses) across the predictive feature space. (B) Matching of sequenced structures with phenotypically similar structures for which the ground truth end-state was captured. (C) Computational strategy to identify biological processes associated with predicted outcomes. (D,E) Volcano Plot showing biological processes at 96h associated with the predicted somitic:neural ratio at 120h, using [Fraction]-T^mCH^ (D) or [Intensity]-T^mCH^ (E) as a proxy. Horizontal dotted line, Padj > 0.05; Vertical dotted lines, NES < -2 or > 2. (F,G) Raincloud plots showing glycolysis module score per [Fraction]-T^mCH^ (F) or [Intensity]-T^mCH^ quartile (G) at 96h. * padj<0.05, ** padj<0.01, *** padj<0.001. (H,I) Volcano Plot showing biological processes in 96h NMPs associated with predicted somitic:neural ratio at 120h, using [Fraction]-T^mCH^ (H) or [Intensity]-T^mCH^ (I) as a proxy. Horizontal dotted line, Padj > 0.05; Vertical dotted lines, NES < -2 or > 2. (J,K) Raincloud plot showing Δ[glycolysis-OxPhos] in NMPs per [Fraction]-T^mCH^ (J) or [Intensity]-T^mCH^ quartile (K) at 96h. (L) Dot plot showing average expression of indicated genes encoding glycolytic pathway genes per [Fraction]-T^mCH^ quartile at 96h in NMPs. (M) Heatmap showing average expression of indicated genes per glycolytic module score quartile. (N) Dot plots showing correlations of expression of indicated genes at 96h in all cells (top) or NMPs (bottom). Dot size scales with *R*.

To identify underlying processes, we correlated pseudo-bulk transcriptome data with dominant predictive features (96h-*[fraction]*-T^mCH^ and 96h-*[mean_intensity]*-T^mCH^). We then ranked genes based on their correlation with either and performed Gene Set Enrichment Analysis (GSEA), which revealed a single prominent hit: “glycolysis” (**Figure 5C-E**). This was corroborated by computing module scores for glycolytic pathway genes, which showed that inferred glycolytic activity is increased in samples predicted to form successful structures (**Figure 5F,G**).

In chick embryos, high glycolytic activity in the posterior end was linked to NMP maintenance and their mesodermal exit.^34^ Indeed, NMPs and mesoderm-biased cells had a higher glycolysis module score than neural-biased cells at 96h (**Figure S6C**). We therefore tested if the observed relationship between inferred glycolytic activity and higher 96h-*[fraction*]-T^mCH^ and/or *[mean_intensity*]-T^mCH^ was conserved in the NMP pool. To this end, we extracted the NMP cluster *in silico*, and performed the same analysis as for all 96h cells. Strikingly, this not only confirmed a positive enrichment for the “glycolysis” term, but additionally revealed a negative enrichment for “oxidative phosphorylation” (OxPhos) (**Figure 5H,I**). Computation of module scores for glycolysis and OxPhos pathway genes provided additional evidence, showing higher glycolysis scores, and lower OxPhos scores, as 96h-*[fraction]*-TmCH or 96h-*[mean_intensity]*-T^mCH^ increased (**Figure 5J,K and S6D,E**). Analysis of the expression of individual genes in the NMP pool showed that for nearly all glycolytic pathway genes expression increased as a function of 96h-*[fraction]*-T^mCH^ or 96h-*[mean_intensity]*-T^mCH^ (**Figure 5L and S6F**). Since the aforementioned study provided evidence for a regulatory FGF-glycolysis-WNT loop in the posterior end of the chick embryo^34,35^, we assessed the relationship between FGF/WNT signaling and glycolysis. We found that higher inferred glycolytic activity was associated with increased expression of FGF/WNT pathway genes (**Figure 5M and S6G,G’**). Moreover, expression of WNTs, FGFs and their downstream targets involved in NMP maintenance and decision making strongly correlated with the expression of genes encoding rate-limiting enzymes of glycolysis (*Pfkl*, *Pkm*, *Pgk1*), as well as *Ldha*, which converts pyruvate to l-lactate (**Figure 5N**). Finally, we found a strong correlation with transcription factors (TFs) governing NMP maintenance and neural vs mesodermal decision making (**Figure 5M,N**).

In sum, these findings show that the metabolic profile of the early NMP pool predicts the capacity of structures to coordinate somite and neural tube formation. Higher glycolytic activity is associated with better outcome, concomitant with increased expression of FGF/WNT pathway genes and their targets. Hence, our data strongly suggest that variable FGF-glycolysis-WNT activity underlies phenotypic variation.

### Divergent metabolic and signaling states upon WNT pulse

Motivated by these findings, we tested if we could detect distinct metabolic and morphogen signatures at earlier stages. To this end, we leveraged the predictive power of 72h-*[mean_intensity]*-T^mCH^ (**Figure 3,4 and S5**), which correlated well with *T* expression (**Figure S6H**). Structures with increased 72h-*[mean_intensity]*-T^mCH^ displayed an increased primitive streak signature at the expense of pluripotency, leading up to an enhanced mesodermal signature 24 hours later (**Figure 6A,B and Figure S6I**).

**Figure 6.**
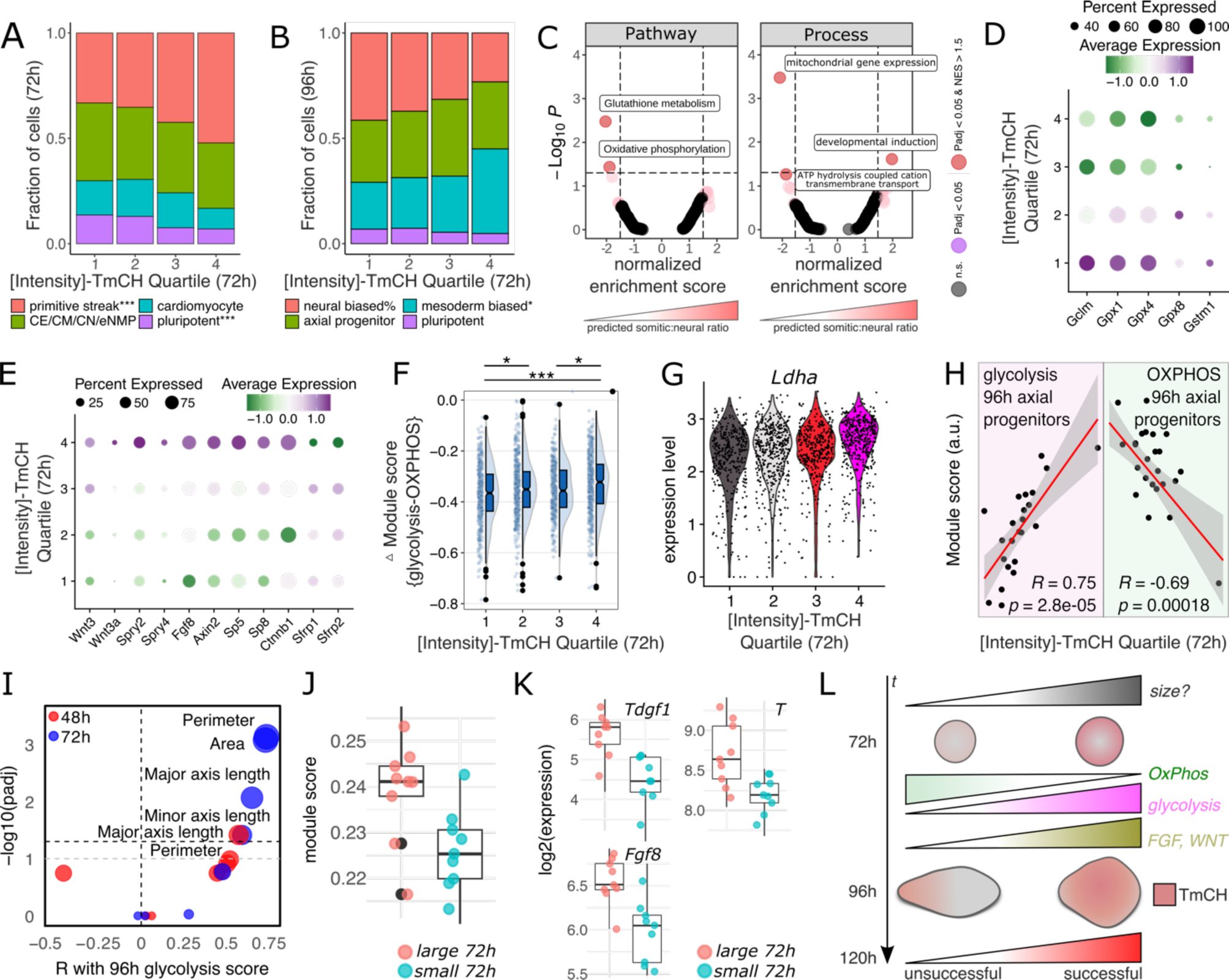
Divergent metabolic and signaling states after WNT pulse. (A,B) Cumulative bar plot showing the fraction of cells in each color-coded annotated state for [Intensity]-T^mCH^ quartiles at 72h (A) and 96h (B). % p=0.05, * p<0.05, *** p<0.001 (ANOVA with BH FDR correction). (C) Volcano Plot showing pathways and processes at 72h associated with a low and high predicted somitic:neural ratio at 120h. Horizontal dotted line, Padj > 0.05; Vertical dotted lines, NES < -1.5 or > 1.5. (D) Dot plot showing average expression of selected genes involved in “Glutathione Metabolism” per [Intensity]-T^mCH^ quartile at 72h. (E) Dot plot showing average expression of selected genes involved in “Developmental Induction” per [Intensity]-T^mCH^ quartile at 72h. (F) Raincloud plot showing Δ[glycolysis-OxPhos] in NMPs per [Fraction]-T^mCH^ (J) or [Intensity]-T^mCH^ quartile (K) at 72h. * padj<0.05, *** padj<0.001. (G) Violin plot showing *Ldha* expression per [Intensity]-T^mCH^ quartile at 72h. (H) Scatter plot showing correlation of [Intensity]-T^mCH^ quartile at 72h and Glycolysis (magenta) or OxPhos (green) module score at 96h. Black dots are individual samples. Grey area indicates 95% CI. (I) Plot showing correlations of bright-field morphometric features at 48h (red) and 72h (blue) with 96h glycolysis module score. Dot size scales with -log10(Padj). (J) Box plot showing the glycolysis module score in large and small structures at 72h (p < 0.05). (K) Box plots showing the log2(expression) of all genes significantly (Padj < 0.05) differentially expressed between large and small structures at 72h. (L) Schematic summarizing the findings.

Lower T^mCH^ levels at 72h were associated with enriched signatures of OxPhos and related processes, including glutathione metabolism - a defense against respiration-induced reactive oxygen species (**Figure 6C,D**). Vice versa, structures with higher T^mCH^ levels showed enrichment of a “developmental induction” module comprising morphogens implicated in NMP maintenance and mesoderm formation (e.g. *Wnt3*, *Wnt3a, Fgf8*), and their downstream targets (**Figure 6C,E**). Notably, the expression of secreted WNT inhibitors *Sfrp1/2* - knockdown of which leads to a multipolar phenotype^36^ - decreased with higher T^mCH^ levels (**Figure 6E**).

Although glycolysis was not among the significantly enriched processes, module score analysis not only confirmed that structures with higher T^mCH^ levels at 72h had lower OxPhos scores, but also revealed higher glycolysis scores and gradually increased *Ldha* expression (**Figure 6F,G and Figure S6J**). Moreover, 72h T^mCH^ levels were predictive of inferred OxPhos and glycolysis activity at 96h (**Figure 6H**). Finally, a targeted analysis showed that an enhanced glycolytic state at 96h correlated with aggregate size at 72h, raising the intriguing possibility that early size differences set the stage for divergent metabolic activity, ultimately impinging on differentiation outcome (**Figure 6I**). Indeed, larger 72h aggregates displayed higher inferred glycolytic activity, concomitant with increased expression of *T*, *Fgf8* and Nodal co-receptor *Tdgf1* (*Cripto*) (**Figure 6J,K**).

Collectively, our data strongly suggest that divergent metabolic and morphogen signaling states drive phenotypic variation. Higher glycolysis-WNT activity results in harmonious formation of somitic and neural tissue, ultimately resulting in successful TLSs (**Figure 6L**).

### Metabolic interventions correct unsuccessful differentiation outcome

The newly identified association between differentiation outcome and divergent metabolic activity in NMPs prompted us to test if the coordinated development of neural and somitic tissue could be tuned by metabolic state alterations. We subjected a total of 966 stembryos to metabolic interventions designed to modulate OxPhos-glycolysis balance and assessed TLS morphology and somitic : neural ratio at 120h (**Figure S7A,D,E**). Specifically, we treated structures from 72-96h or from 96-120h with the mitochondrial complex I inhibitor rotenone^37^ or 2-Deoxy-D-glucose (2-DG), a compound that competes with glucose to bind hexokinase, the first rate-limiting enzyme of glycolysis^38,39^ (**Figure 7A and S7A**).

**Figure 7.**
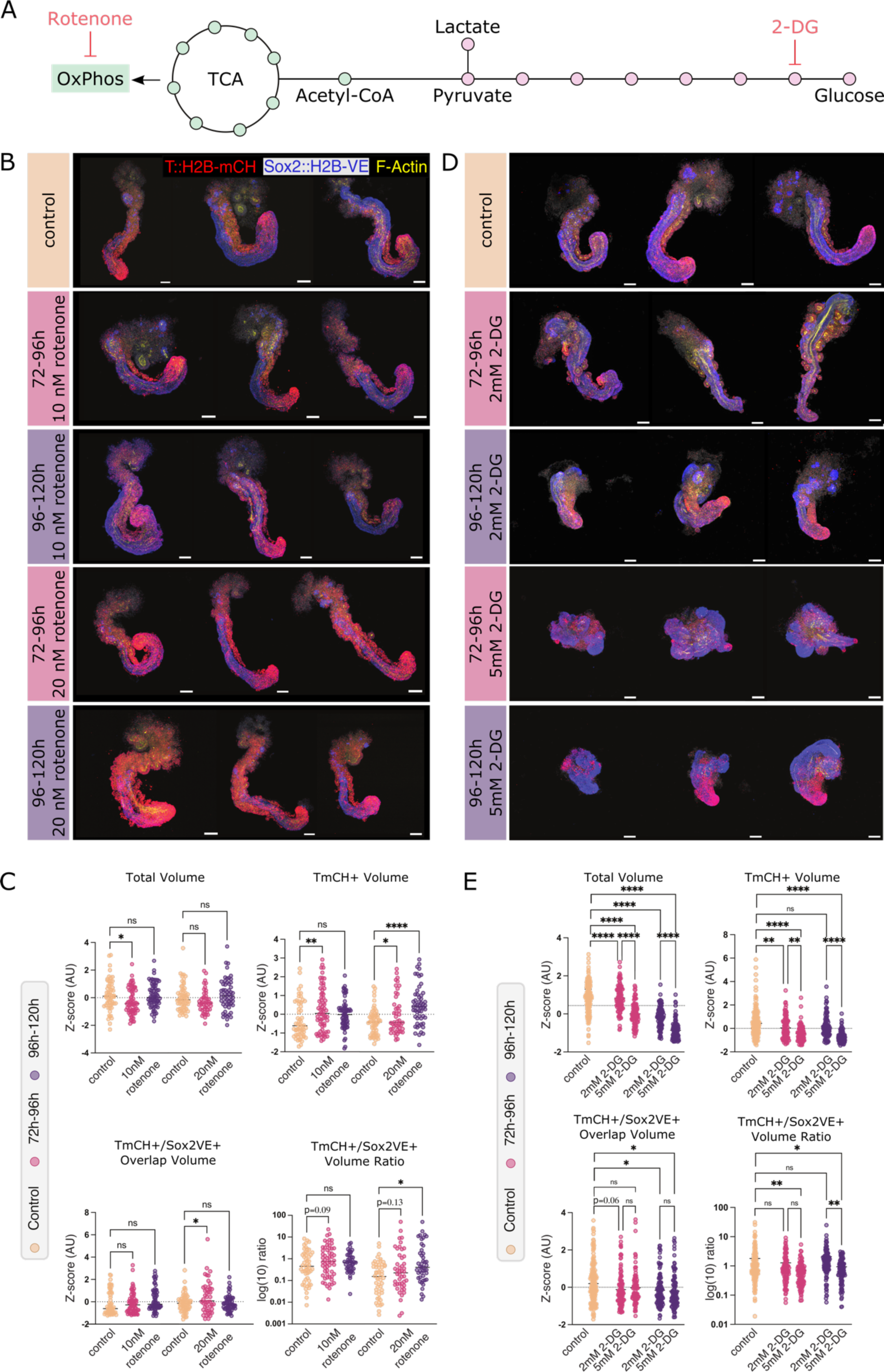
Metabolic interventions correct derailed differentiation outcome. (A) Schematic representation of central carbon metabolism. The metabolic flux of glucose can be modulated using different inhibitors. Inhibitors in red designed to alter OxPhos-glycolysis balance (see text). For a detailed schematic see Figure S7A. (B) Representative 3D maximum intensity projections (3D-MIPs) of 120h control and Rotenone-treated TLSs. Scale bars, 100 µm. (C) Quantifications of 120h TLSs treated with Rotenone 10nM (72h-96h, n=69; 96h-120h, n=53) or 20nM (72h-96h, n=46; 96-120h, n=55), and their respective controls (Control for 10nM, n=59, Control for 20nM, n=63). *, p<0.05, **, p<0.01, ***, p<0.001, ****, p<0.0001. (D) Representative 3D-MIPs of 120h control or 2DG-treated TLSs. Scale bars, 100 µm. (E) Quantifications of 120h TLSs treated with 2-DG 2mM (72h-96h, n=100; 96h-120h, n=105) or 5mM (72h-96h, n=100; 96-120h, n=65), compared to control (n=220). *, p<0.05, **, p<0.01, ***, p<0.001, ****, p<0.0001.

Treatment with rotenone from 96-120h did not drastically affect overall morphology, but clearly increased the T^mCH+^ volume (both absolute and fraction), concomitant with increased somite formation at the highest concentration (**Figure 7B,C and Figure S7B,D,F**). Vice versa, Sox2^VE+^ volume fraction was reduced (**Figure S7F**). Treatment from 72-96h increased not only T^mCH+^ volume, but also the volume with tissue-level co-localization of T^mCH^ and Sox2^VE^ at both 96h and 120h at the highest rotenone concentration (**Figure 7B,C and Figure S7G,H**). Altogether, these results show that a metabolic intervention that induces glycolysis at the expense of OxPhos leads to more balanced lineage allocation of NMPs - also evident from the increased T^mCH+^/Sox2^VE+^ volume ratio (**Figure 7C**) - and that early, but not late, enhancement of glycolysis expands the NMP pool (**Figure 7C and S7G,H**).

In contrast to rotenone, 2-DG treatment resulted in pronounced alterations in overall morphology, showing a dose-dependent reduction of total volume and elongation (**Figure 7D,E and S7C**). At the highest concentration, TLS morphology was severely disrupted with aberrant formation of disorganized neural tissue and severely hampered somite formation (**Figure 7E and S7I**), emphasized by quantitative analysis demonstrating a decreased T^mCH+^/Sox2^VE+^ volume ratio and somite number (**Figure 7E and S7J**). Although the lower concentration did not quantitatively affect the T^mCH+^/Sox2^VE+^ volume ratio (**Figure 7E**)), clear defects were observed. Treatment from 72-96h resulted in disorganization of (posterior) somitic tissue with “scattering” of somites (**Figure 7D and S7I**), whereas treatment from 96-120h completely abolished segmentation of somitic (T^mCH+^) tissue (**Figure 7D and S7I,J**). Finally, quantification of the T^mCH+^/Sox2^VE+^ overlapping volume showed that 2-DG decreased the size of the NMP pool (**Figure 7E**). In sum, effects of 2-DG are dose- and time-dependent; high concentration increases neural at the expense of somitic tissue and inhibits elongation, low concentration affects segmentation. Both concentrations decrease the size of the NMP pool.

Collectively, the metabolic interventions show that the phenotypic landscape of the stembryo can be tuned by altering the balance between oxidative phosphorylation and glycolysis. Most importantly, favoring glycolysis can correct neural lineage bias. Furthermore, the data suggest the presence of an early temporal window in which enhanced glycolysis increases the size of the NMP pool.

## DISCUSSION

Viktor Hamburger famously stated that “the embryo is […] always right”.^40^ In contrast, stembryos frequently deviate from this ideal state, manifesting as phenotypic variability. Such seemingly stochastic variation represents a major hurdle. Here, we demonstrated that biological processes underlying phenotypic variation can be identified through integrated molecular-phenotypic profiling.

We focused on the variable propensity to form somites and a neural tube from an NMP pool. Our finding that this divergent propensity can be detected prior to the end-state and associates with variable WNT signaling activity aligns with recent work in gastruloids.^41^ Our unique integrative strategy in combination with functional experiments strongly suggest that this divergence is driven by variable metabolic activity, and can be controlled by metabolic interventions. Specifically, we reveal that glycolytic versus OxPhos activity regulates NMP decision making and, potentially, maintenance, presumably by regulating FGF and/or WNT signaling activity.

Consistent with our findings, a parallel preprint showed that decreasing glucose concentrations in early gastruloids progressively shifts germ layer proportions away from mesodermal derivatives towards more neuroectodermal cell fates.^42^ Furthermore, the authors show that the addition of Nodal or Wnt signaling agonists can rescue mesoderm and endoderm development in glycolysis-inhibited gastruloids, suggesting that glycolytic activity is an important activator of signaling.^42^ Hence, this and our work provides further support for the idea that metabolic pathways can act as regulators of patterning and morphogenesis during embryogenesis. The concept of metabolic signaling, rooted in the metabolic gradient theory^43^, has picked up interest.^44^ Some work focused on gastrulating amniote embryos, and is therefore of particular interest in light of our discoveries.

In chick, suppressing glycolysis alters NMP differentiation potential, promoting a shift towards neural at the expense of mesodermal differentiation.^34^ An FGF-glycolysis-WNT loop was proposed, where an FGF signaling gradient establishes posterior-to-anterior glycolytic activity, leading to a corresponding gradient in WNT signaling.^34,35^ Our findings suggest that this loop operates in stembryos and that its variable activity underlies phenotypic variation.

On the contrary, in mouse embryo explants glycolysis was reported to negatively regulate WNT signalling.^38,45^ Although the authors suggested that the discrepancy could be rooted in short-term negative^38^ versus long-term positive regulation^34^, our data show that - at the same time-point - higher glycolytic activity is positively associated with expression of WNTs and their targets. Our discovery that the FGF/WNT/glycolysis connection is conserved in the scalable and tractable stembryos presents an exciting opportunity to unravel the intricate spatiotemporal dynamics of feedback.

Pyruvate produced by glycolysis can either enter the citric acid cycle and be catabolized by OxPhos, or converted into lactate by Ldha. The latter phenomenon, known as aerobic glycolysis or the Warburg effect, results in lower ATP yields.^46^ In chick, aerobic glycolysis activity is higher in NMPs and their mesodermal derivatives.^34^ The resulting lactate production establishes a pH gradient, favoring the mesodermal over neural lineage choice, primarily through the activation of WNT signaling.^35^ *In vitro* PSM differentiation tied this to non-enzymatic β-catenin acetylation. Such regulation downstream of β-catenin stabilization might explain the observed inter-structure variation in WNT signaling pathway activity despite an initial pulse with WNT activator CHIR (an inhibitor of GSK3-β which phosphorylates β-catenin and targets it for degradation^47^).

During the embryonic stages modeled here, the embryo develops under hypoxia (1.5-8%^48^). One aspect of the hypoxic response involves a transition in cellular metabolism, from OxPhos to glycolysis.^49^ We recently reported that in hypoxic gastruloids the relative fraction of NMPs and their somitic progeny is increased.^50^ Hence, a hypoxia-induced shift from OxPhos to glycolysis could be a physiological means to balance neural versus mesodermal decision-making of NMPs. We speculate that small variations in size could set the stage for such divergent metabolic activity.

Recent work investigated the role of glucose metabolism during mouse peri-gastrulation stages using (st)embryos.^51,39^ One study showed that during early gastrulation (∼E6.5 - ∼E7.25) mesoderm fate acquisition and maintenance is governed by glucose metabolism via the Hexosamine Biosynthetic Pathway (HBP), but mesoderm migration and lateral expansion requires glycolysis.^51^ Complementary work in gastruloids suggested that mannose, but not glycolysis, is crucial for the early induction of *T*.^39^ While at first glance some of these findings seem at odds with our discoveries, most of the effects we report here relate to later developmental stages (embryo equivalent: ∼E7.5-E9^4^), during which mesoderm and neural cells are specified from a pool of NMPs. Hence, our data could reflect the idea that late-stage glycolysis mostly becomes relevant after (neuro)mesoderm specification from the primitive streak. Alternatively, or additionally, lactate induced mesoderm specification (via regulation of WNT signaling) could act on top of mannose metabolism.^39^

An intriguing phenomenon in a subset of structures comprised balanced somitic and neural tissue formation in an “inside-out” configuration, with neural tubes enveloping non-segmented somitic tissue. This uncoupling of cell state composition and morphogenesis challenges the classification process based on end-state BF images, resulting in substantial overlap of end-states in the PLSR latent spaces, ultimately impinging on accuracy scores. Further advancing the framework by incorporating longitudinal confocal instead of wide-field imaging, thereby extending the feature space with e.g. detailed spatial distribution, progenitor field appositions, and cellular geometry, could significantly enhance predictive capabilities. However, this will pose challenges related to throughput, phototoxicity, and real-time predictions and interventions.

Finally, our study underscores that while integrated molecular-phenotypic profiling is essential for identifying processes driving phenotypic variation, machine learning applied to sparse longitudinal imaging data suffices to predict phenotypic end-states. This is further emphasized by a recent preprint employing machine learning to predict endoderm end-state morphology.^52^ Such predictive power can be leveraged to discover design principles that govern developmental canalization (as we show here), and enhances the potential of stembryos for applications that require high reproducibility, including disease modeling and reproductive toxicity studies. For example, our strategy is useful to stratify samples across experimental conditions to ensure a balanced representation of predicted outcomes, thereby reducing false positive and false negative rates. Moreover, the possibility to obtain (semi-)paired measurements (by subjecting overlapping samples in the predictive state space to different treatments) could reveal state-specific effects.

We employed the integrated molecular-phenotypic profiling to identify biological processes underlying (phenotypic variation in) coordinated formation of somitic and neural tissue. However, our dataset and strategy can be (re)-employed to investigate the roots of many (variable) outcome measures, e.g. elongation or curvature. While these will comprise correlations, the high throughput and potential for global and personalized interventions make stembryos an excellent system for investigating causality, dissecting feedback loops, and, ultimately, controlling shape and patterning.

## METHODS

### Mouse Embryonic Stem Cell Culture

Mouse Embryonic Stem Cells (mESCs) carrying a T::H2B-mCherry and Sox2::H2B-Venus reporter^10^ were cultured under standard Serum+LIF+feeder conditions (400 mL Knockout Dulbecco’s Modified Eagle’s Medium (DMEM) (4500 mg/ml glucose, w/o sodium pyruvate) (Gibco) supplemented with 75 mL ES cell tested FCS, 5 ml 100x L-glutamine (200 nM) (Lonza #BE17-605E), 5 mL 100x penicillin (5000 U/ml)/ streptomycin (5000 μg/ml), 5 ml 100x non-essential amino acids (Gibco #11140-35), 1 mL 500x β-mercaptoethanol (5mM, 1000x Invitrogen), 5 mL 100x nucleosides (Chemicon) and 5µl 10000x of Murine Leukemia Inhibitory Factor (LIF) (Sigma-Aldrich, #ESG1107) on 6cm plates (Sarstedt, # 83.3901.300) coated with 0.1% gelatin and a layer of mitotically inactive fibroblasts. Culture medium was refreshed every 24h and mESCs were split using Trypin-EDTA (Thermofisher, #ESG1107) in a dilution of 1:10 every 48h. The cells were passaged twice before proceeding to TLS formation.

### Generation of Gastruloids & Trunk-Like-Structures

Gastruloids and TLSs for were generated following the previously published protocol.^10,15^ In short, mESCs were feeder-freed by sequential plating of the cells on 0.1% gelatin-coated 6-well plates and incubating the cells at 37°C for 25’, 20’ and 15’ respectively. Cells were then washed with 5mL pre-warmed PBS supplemented with MgCl_2_ + NaCl (PBS+/+) (Sigma, #D8662-500mL), spun down for 5’ at 300g, followed by a second wash with 5mL of pre-warmed NDiff227 medium (Takara). After pelleting (5’, 300g) and gentle trituration in 500µl NDiff227, cells were counted using an automatic cell counter (Invitrogen™ Countess™ 3 FL Automated Cell Counter) and diluted in NDiff227 to a final concentration 10 cells/µl. 30µl of the cell suspension (∼300 cells) were then plated into individual wells of a ultra-low attachment U-bottom 96 well plate (Costar, #7007) using a multichannel pipette, and placed in an incubator (37°C, 5% CO_2_). After 48h, the aggregates were pulsed with 3μM CHIR99021 (CHIR) (Tocris, #4423) diluted in 150µl NDiff227 per sample. At 72h, 150µl medium was refreshed with new pre-incubated NDiff227. Between 92h-96h (96h for all samples used for integrated molecular-phenotypic profiling), TLS formation was initiated by supplementing the gastruloids with 5% (final v/v) Growth-factor-Reduced Matrigel (Corning #356231) diluted in NDiff227 medium.

In the case of the rotenone and 2-DG treatment, inhibitors were added for 24h from 72h to 96h or 96h to 120h. The drugs were diluted in the Ndiff227. In the case of rotenone, a 1mM working stock was generated by diluting rotenone (Sigma-Aldrich, #R8875-1G) in DMSO (Sigma-Aldrich, #41639-100ML). The calculated volume of 1mM rotenone solution was then mixed with NDiff227 media to reach the final concentration of 10nM and 20nM. The equivalent volume of DMSO was added to the control. Similarly, 2-DG (Sigma-Aldrich, #D8375-1G) was weighted using a precision balance and diluted in the corresponding volume of NDiff227 to reach the desired concentration of 2mM and 5mM.

### Whole-mount Immunofluorescence (WIFC)

Structures were harvested using a wide-boar p200 pipette tip and transferred to an 8-well glass-bottom plate (Ibidi #80827). The structures were then washed three times with PBS++/BSA (PBS supplemented with MgCl2 and CaCl2 (Sigma-Aldrich, #D8662-6X500ML) supplemented with 0.5% BSA) and three times with PBS. Thereafter the structures were fixed with 4% PFA for 1h at 4°C, followed by 3 washes with PBS. The structures were then permeabilized with PBS-T (PBS supplemented with 0.05% (v/v) Triton-X) and subsequently blocked overnight with blocking solution (PBS supplemented with 0.05% (v/v) Triton-X and 10% FCS). TLSs were then incubated with the primary antibodies at a dilution of 1:250 for 72h to 96h. Subsequently, structures were washed three times with PBST-X and left in the Blocking solution overnight. The next day, the secondary antibody at a dilution of 1:500 was added and incubated for 24h. TLSs were then washed three times with blocking solution, once with PBST-X and then post-fixed with 4% PFA for 20’ to 1h. Then, samples were washed with 0.02M Phosphate Buffer (PB) (25mM NaH2PO4 + 75mM Na2HPO4, pH 7.4) twice and embedded in a drop of low-melting agarose to stabilize them for imaging. Finally, the structures were cleared using RIMS (133% (w/v) Histodenz (Sigma-Aldrich, #D2138) diluted in 0.02M Phosphate Buffer supplemented with 0.01% Tween) overnight. A list of all the antibodies used in this manuscript can be found in **Table S2**.

### Imaging

#### Wide-field Imaging

Widefield imaging of live samples was performed using Zeiss CellDiscoverer7 (CD7) at 48h, 72h, 96h and 120h. The temperature and CO_2_ levels were kept constant at 37°C and 5% respectively for the entire duration of the imaging. The samples were imaged *in situ*, in a 96-well ultra-low attachment U-bottom plates using a Plan-Apochromat 5×/0.35 objective, a TL LED Lamp and LED-Module 590nm light source with adequate emission filters. The images were acquired at 8-bit depth and the image was scaled to 101.36mm × 65.35mm.

#### Spinning Disk Imaging

Fixed and stained structures were imaged on a spinning disc microscope (Olympus IXplore SpinSR using Yokogawa W1 with Borealis laser source, iXon Ultra 888 camera, Olympus 10x/0.4 U Plan SApo Air objective and appropriated laser/filters for 405, 488, 568 and 640nm emission fluorophores). A Z-stack of approx. 250μm was taken with a z-step of 9μm and all images were acquired with a pixel size of 0.65×0.65μm^2^.

### Imaging processing and analysis

#### Manual classification of TLSs based on bright-field imaging

Criteria for classification were the same as applied in Veenvliet et al., 2020.^10^ Structures were classified as “successful” when at least four neighboring segments had developed along the antero-posterior axis at 120h. Segments were defined as sub-structures that displayed i) indentations, and ii) opposite curvatures at segment borders. Structures that displayed two or more axes were classified as “multiple axes” (multipolar), others as “one axes” (unipolar).

#### Confocal imaging processing

For confocal image processing, multi-channels and multi-slice images were used. Adjustments to the channels’ colors and intensities and contrast enhancement for the thumbnails were made based on the user. A user-drawn initial crop box is repeated by FIJI for all subsequent structures, while the suggested position of the crop box can be optionally readjusted by the user. Channels of interest can be selected for creating a 3D Projection (3DP) of the image. The cropped z-stack is saved as TIFF only, while the resulting projections and each projected individual channel (in grayscale) are saved as TIFF and JPEG files. Finally, a scale bar of 100μm is applied to the final images. All image processing was executed using Fiji v1.54f.

#### Mean Intensity quantification

For quantifying volume fractions, multi-channel and multi-slice images were used. The images were processed with background subtraction of the modal value for each slice in the image stack to remove background noise. For each channel, the thresholds are set based on the user-provided parameters for lower and upper thresholds. The appropriate channel is then renamed according to the user-provided channel names. Particles (regions of interest) are analyzed in the image using the Analyze Particles function and a summary of the z-stack’s results is generated. The results are then saved as a CSV file in the specified output directory, with a filename based on the channel name and, if applicable, the condition name. All image analysis was executed using Fiji v1.54f.

#### Volume and Overlap Volume Fraction quantification

For quantifying overlap volume fractions, multi-channel and multi-slice images were used. Bio-Formats Importer was utilized to open the image files and process them as grayscale hyperstacks. The images were processed with background subtraction of the modal value for each slice in the image stack to remove background noise. The appropriate channel number, lower threshold and name is provided for four different channels (A, B, C, and D) (NOTE: the lower threshold was kept unchanged with respect to the one used to quantify intensities). The JACoP plugin^53^ performs colocalization analysis. Channels can be combinatorially compared, allowing users to assess colocalization and interaction patterns between different channels in the microscopy images. The results of each channel comparison are saved as separate results files (Results.csv) in the specified output directory. All image analysis was executed using Fiji v1.54f

#### Data processing and Statistics

The Sox^VE+^, T^mCH+^ and Sox^VE+^ and T^mCH+^ Overlap Volume were calculated by computing the sum of the area of each individual channel over the total amount of z-stacks of a structure. The Volume Fraction was calculated by dividing the Volume of each individual channel by the total Volume of the respective structure (DAPI Volume). The T^mCH+^ and Sox^VE+^ Ratio was computed by dividing the T^mCH+^ volume by the respective Sox^VE+^ per each structure. Z-scores were calculated across all conditions on a per experiment basis using the formula:

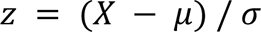

where *X* is the sample value, μ is the population mean, and σ is the standard deviation.

The statistical analysis was carried out using commercial software GraphPad Prism (v. 9.5.1 (528)). The following statistical tests were performed:

Figure 1C: Unpaired two-tailed t-test

Figure 3E-J: Two-way ANOVA with Tukey HSD test for multiple comparisons

Figure 5F,G,J,K, Figure 6F: One-way ANOVA with Tukey HSD test for multiple comparisons

Figure 7C,E: One-way ANOVA, Dunnet multiple comparison correction.

Figure S1C,D: Unpaired two-tailed T-test.

Figure S1F: Two-way ANOVA, Tukey’s multiple comparison correction.

Figure S5D: Two-way ANOVA with Tukey HSD test for multiple comparisons

Figure S6D,E: one-way ANOVA with Tukey HSD test for multiple comparisons

Figure S7F,H: One-way ANOVA, Dunnet’s multiple comparison correction.

Figure S7J: Two-way ANOVA, Tukey’s Multiple comparison correction.

#### Somite quantifications

The number of somites was quantified in 20 random selected images per condition manually using the pre-installed Cell Counter plug-in in Fiji v1.54f. The criteria to consider a structure as somite was: 1) the structures need to be round, 2) show a single internal lumen defined by phalloidin staining, 3) be T^mCH+^ and Sox2^VE-^. Only when all three criterias were met the structure was counted as somite. We did not threshold the somitic structures according to size.

#### Axis Length quantification

To determine the axis length of structures images using volumetric imaging, the volumetric images were projected to 2D using a mean projection. These images were then segmented using a pipeline consisting of denoising (Gaussian blur), thresholding (Otsu Method^54^) and a binary fill holes operation (all implemented in scikit-image^21^). The segmented structures were then skeletonized using the skeletonization operation implemented in scikit-image. These topological skeletons were further processed using ToSkA^55^ to determine the length of the longest path within the topological skeleton, a proxy for axis length.

#### Image Analysis of Widefield Microscopy Images

To extract focused images from multi-focus image stacks, we employed the Gaussian-based stack focuser implemented in Fiji^19^ for both the bright-field and T^mCH^-Channels. The bright-field (BF) images underwent min-max scaling and were subsequently segmented using Cellpose 2^56^. Segmentation involved the use of two custom-trained models: The cyto2-model, trained iteratively on 90 manually corrected 96h masks, was employed for segmenting the 72h and 96h images, while the cyto2-model trained on 27 manually corrected 48h masks was used for segmenting the 48h images. This resulted in the BF masks. For the segmentation of the TmCH domain, we used scikit-image^21^. Initially, a global threshold was determined using the Otsu method^54^ for all 96h images. Subsequently, the images underwent denoising via Gaussian blur before being thresholded using the global threshold. Lastly, any T^mCH^ domain regions outside of the BF mask were removed, resulting in the final T^mCH^ domain masks. To ensure that each mask image contained only one object, a filtering step was implemented to retain only the object with an area closest to the median area of all objects.

To expand the feature space, we utilized several feature extraction methods. Firstly, simple shape features of the BF masks were calculated using scikit-image^21^. Secondly, the BF masks were straightened using MOrgAna^17^, to remove any influence of curved structures and subsequently more complex shape features implemented in MOrgAna were calculated, resulting in straightened shape features, which have the prefix: str. The image straightening implemented in MOrgAna was also used to determine how polarized the T^mCH^ domain was along the major and minor axis. For this a custom feature extraction method was used, that straightens the T^mCH^ image, alongside the BF mask and determines the distance between the BF mask centroid and the TmCH domain centroid along both the major and minor axes of the straightened BF mask. Lastly, to calculate T^mCH^ features cellprofiler^57^ was used, as it not only can measure intensity, but also intensity distribution features. For this both the bright-field mask and the T^mCH^ domain mask were used. Intensity Features measured inside the T^mCH^ domain mask have the suffix T^mCH(mCH)^ and intensity features measured in the BF mask have the suffix T^mCH(BF)^. Additionally, cellprofiler was used to measure simple shape features of the T^mCH^ domain mask, which also have the suffix T^mCH^. An overview of the measured features can be found in Supplementary Note 1.

### MULTI-Seq sample preparation

The multiplexed single-cell experiment was performed as previously described.^23,50^ Briefly, structures were generated as described above. Following wide-field imaging, 72 structures at 48h, 48 at 72h, 24 at 96h and 24 at 120h were picked with a p200 cut tip and transferred in a new well of a 96-well plate. Next, structures were washed twice with ice cold PBS and trypsinized in 20µl TrypLE Express (*Gibco*) for 25 minutes in the incubator at 37°C, pipetting every 5 minutes to help dissociation. Multi-seq labeling was then performed as previously described.^23^ Briefly, single-cell suspensions of each sample were incubated with a unique BC-Lipid modified oligonucleotide “anchor” mix (200nM each final) 5 minutes on ice. Next, a 200nM “co-anchor” mix was added to each sample and cells were incubated for an additional 5 minutes on ice. The reaction was then quenched by addition of 200µl 1xPBS/1%BSA, and cell suspensions were then washed twice with 1xPBS/1%BSA in the plate. Next, all samples for each day were pooled in a 1.5 ml DNA lowBind tube in 1xPBS/1%BSA, and cells pelleted by centrifugation. The recovered cells were resuspended in 1xPBS/0.4%BSA, counted and then subjected to 10x single-cell RNA-seq using the 10x Genomics Chromium™ Single Cell 3’ v3.1 (one reaction for gastruloid pool at 48, 72, and 96h; two reactions for TLS pool at 120h) (see section below for 10x processing). MULTI-seq oligonucleotides’ sequences are listed in **Table S3**.

### Single-cell RNA sequencing

Following MULTI-seq barcode labeling, single-cell RNA-seq (scRNA-seq) experiments were performed as previously described^23,50^. Briefly, single-cell suspensions were subjected to GEMs encapsulation and libraries generated according to the manual to recover the cDNA fraction. Quality and concentration of the obtained libraries were measured using Agilent High Sensitivity D5000 ScreenTape on an Agilent 4150 TapeStation. Libraries were sequenced with a minimum of 400 million paired end fragments according to parameters described in the manual. To recover the Multi-seq barcodes, two modifications were introduced in the standard workflow: (i) during the cDNA amplification step 1µl of an oligonucleotide to enrich for the MULTI-seq BCs was added to the reaction and (ii) after cDNA amplification and incubation with SPRIselect beads, the Multi-seq BCs containing supernatant was collected and subjected to further SPRIselect beads incubation in order to recover the Multi-seq BCs as previously described.^23^ Multi-seq BCs recovery and integrity were measured using Agilent High Sensitivity D5000 ScreenTape on an Agilent 4150 TapeStation. The obtained material was then used as input for Multi-seq BCs library preparation (see section below).

### Multi-seq barcodes library preparation from scRNA-seq

Multi-seq BCs libraries were prepared as previously described. ^23^ Briefly, 10ng input material obtained from the 10x cDNA purification (see section above) was used to perform library PCR using KAPA HiFi HotStart ReadyMix (*Roche, KK2601*) in 50µl reaction with the following steps: 95°C 5 minutes/ 98°C 15sec – 60°C 30sec – 72°C 30sec (13 cycles)/ 72°C 1 minutes/ 4°C hold. Next, AMPure XP beads (*Beckman, A63881*) cleanup (1.6X) was performed to purify the Multi-seq BC libraries. Quality and concentration of the obtained libraries were measured using Agilent High Sensitivity D5000 ScreenTape on an Agilent 4150 TapeStation. Libraries were then sequenced using asymmetric end sequencing (150 cycles kit; 28/91 FC-410-1002) on a Novaseq platform at a minimum of 50 million fragments per sample. Oligonucleotides sequences used for library preparations are listed in Table S3.

### Computational Analysis

#### Pre-processing and demultiplexing

Cell Ranger pipeline version 3 (10x Genomics) was used to de-multiplex the raw base call files, generate fastq files, map to the mouse reference genome mm10, filter the alignment and count barcodes and unique molecular identifiers. To de-multiplex samples within our single MULTI-Seq scRNA-seq dataset, we used the deMULTIplex R package (version 1.0.2).^23^ In short, sample IDs were assigned to cells. Cells with no associated sample barcode (unassigned (unable to provide a sample identity using the deMULTIplex algorithm) and negative (cells without barcode), as well as cells with more than one barcode (doublets) were removed for downstream analysis (**Figure S3A,B**).

#### Quality Control (QC), clustering and cell type annotations

After preprocessing, the two scRNA-seq runs for time-point 120h were merged into one single object. Initial QC was performed with Seurat (version 4.3.0).^58^ Single-cell data generated were loaded with a minimum requirement of 3 cells and 200 features (default parameters). Subsequently, cells with unique feature counts < 2,000 & > 80,000 (48h and 120h); < 500 & > 80,000 (72h and 96h) and a mitochondrial fraction above 5% were discarded from the analysis. The Seurat objects after QC were then used for cluster determination. This procedure was performed individually per each time-point. Subsequently, the expression data were independently normalized, and variable features within the three datasets were detected, log-normalized and scaled to 10,000 (default settings). A list of cell cycle markers was used to score for cell cycle stage and to subsequently scale the data with regression out (var.to.regress) of S and G2M phase-related genes^59^. For downstream analysis and visualization of the datasets, a PCA followed by a UMAP (dims=1:30, n.neighbors=10) were run, and clusters were identified using the Louvain algorithm. Next, we manually inspected the distribution of UMI counts, total RNA counts, mitochondrial and ribosomal fraction for each cluster at individual time-points. Based on this additional QC step we removed spurious clusters. Specifically, for 48h, two clusters (one with low n cells (<10) & one with low mitochondrial fraction) were removed, followed by reclustering; for 96h a small cluster was removed with a low mitochondrial and high ribosomal fraction, followed by reclustering; for 120h, a putative doublet cluster (shown by almost double amount of total RNAs counts and UMIs) was removed. The remaining clusters were then further annotated using a combination of classifiers and manual curation to ensure most accurate and reliable annotation of cell states. While for 48h, 96h and 120h the clusters identified by the Louvain algorithm were readily interpretable in terms of marker genes, this proved more challenging for the 72h samples. We therefore first used a publicly available mouse reference atlases (E6.5 to E8.5) to predict the in vivo cell ID using scmap^26,60^. In short, a table of the top variable features across the reference dataset was obtained, and relevant markers were chosen with a criterion of passing the threshold of log fold change of 0.5, and a p-value of 1e-5. Following the curation of the relevant marker table, the top 200 markers were chosen in each reference cluster to train a classifier. The query dataset was then converted to an sce (SingleCellExperiment) object, and the classifier was applied to the query atlas – after filtering to include only the relevant stages (E6.5 and E7.5). Finally, the 72h object was re-clustered using a list of the top 20 marker genes (obtained using the FindAllMarkers function in Seurat with default settings, except for pseudocount.use=0.1) for all in vivo cell types identified by the classifier (FindNeighbors, dims=1:10 and FindClusters, resolution=0.4). To subcluster the 120h cluster with a pluripotency signature, the cluster was subsetted and re-clustered (FindNeighbors, dims=1:10 and FindClusters, resolution=0.05). Gene-sets used to calculate EPCs and PGCLCs module scores were *[Upp1, Klf4, Klf9, Tbx3, Zfp42, Dnmt3l]* and *[Rhox6, Rhox9, Dnd1, Prdm1, BC048679]* respectively.

#### Pseudotime inference

Pseudotime inference of Seurat objects was conducted using the Monocle 3 package (version 1.3.1).^61^ Initially, Seurat objects were converted into cell data sets (cds) within the Monocle framework. Subsequently, cells were clustered using a time-point specific resolution, and a trajectory graph was learned to capture the temporal relationships among cells. Cells were ordered along the trajectory with specified root cells (based on cluster annotations) to establish a pseudo-time continuum. To analyze pseudotime distribution per sample, we integrated Monocle-derived pseudotime values into the metadata of the Seurat object. We then filtered the samples by their occurrence frequency, selecting only those with more than 20 cells. Plotting was then conducted using ggplot2.

#### Calculation of *CV2* for cell type composition

First, we filtered the samples by their occurrence frequency, selecting only those with more than 20 cells. Subsequently, we calculated cell type proportions based on the Seurat clusters. To quantify the variability within each cell state, we computed the squared coefficient of variation (*CV2*) for each cluster:

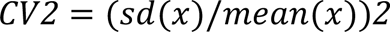

where *sd(x)* is the standard deviation of the values in *x*, and *mean(x)* is the mean of the values in *x*. For the merged_cluster plot, we merged the three clusters with a neural signature.

#### Identification of genes, processes and pathways with high inter-embryo variation

For each timepoint, single-cell RNA-Seq data were pseudo-bulked on a per-structure basis, and pseudo-bulk data were normalized to counts per million (CPM). Genes with no expression in any sample were filtered out, followed by the removal of genes with low mean expression (< 0.1) across all samples. After computing the *CV*^2^, a generalized linear model was fitted to the data to capture the relationship between mean and variance. Highly variable genes were identified based on this model. Subsequently, a winsorization process was applied to mitigate extreme values. Most variable genes were then recalculated using the winsorized data. Finally, a chi-squared goodness-of-fit test followed by FDR correction was applied to assess whether the variability in gene expression across samples is statistically significant (defined as *Padj* < 1e-3).

#### Visualization and statistical comparison of cellular composition

For visualization in the form of stacked boxplots, we used the dittoSeq package (version 1.6.0).^62^ To test for statistically significant differences in cell type proportions between groups, we employed the propeller function as part of the speckle package (version 0.03) using default settings (ANOVA with BH FDR correction).^63^

#### Computation of module scores

To compute somitic and neural module scores, we employed the FindMarkers function in Seurat to find markers specific to the somitic and neural cluster (pseudocount.use=0.1, min.pct.diff=0.25). Next, we extracted the top 25 markers based on log-fold change values, and computed module scores using the AddModuleScore function in Seurat with default settings.

To compute glycolysis and oxidative phosphorylation(OxPhos), gene sets associated with glycolysis and OxPhos were extracted from Malkowska et al., 2022.^64^ Module scores were then calculated for each cell using the AddModuleScore function in Seurat with default settings. To compute NMP-specific scores, we subsetted the 96h Seurat object to only contain the NMP cluster.

#### Correlating features with module scores

Features were added as metadata to the Seurat object. After computation of somitic and neural module scores, we aggregated scores based on a per-structure basis. We then calculated Pearson correlations of these neural and somitic module scores with the complete feature space (after removal of features with a standard deviation of zero) using the *cor* function in the *stats* package. To obtain confidence intervals and p-values for correlations, we used the *corr.test* function in the *psych* package (version 2.3.3). Significant correlations were defined as *Padj* (holm) < 0.05 and *R* > 0.7

#### Correlating features with gene expression

To compute correlations of features of interest with gene expression, single-cell RNA-Seq data were pseudo-bulked on a per-structure basis, and pseudo-bulk data were normalized to counts per million (CPM). We then computed Spearman correlations using the *corr.test* function in the *psych* package.

#### Quartile binning

For quartile binning of samples based on features of interest, we employed the quantile function in R to calculate quantiles at five equidistant levels (0%, 25%, 50%, 75%, and 100%).

#### Differential Expression Analysis (DEG) on pseudo-bulk transcriptomes

First, we filtered the samples by their occurrence frequency, selecting only those with more than 20 cells. Subsequently, we pseudo-bulked on a per-structure basis, and pseudo-bulk data were normalized to counts per million (CPM). Next, we filtered and classified structures as large or small based on their Area (large, > 67,500; small, < 62,500). We then used EdgeR^65^ (version 3.36.0) to compute DEGs. A DGEList object was constructed, and differential expression analysis was performed using the edgeR package. The design matrix was established for a simple comparison between “small” and “large” groups, and dispersion estimation and negative binomial model fitting were executed. Genes with an FDR-corrected p-value < 0.05 were considered significant.

#### Gene Set Enrichment Analysis (GSEA)

For GSEA, we used the WebGestaltR package.^66^ Ranked gene lists based on descending correlation values with the feature of interest were provided as input. Analysis was conducted for both Biological Processes and Molecular Functions from gene ontology, as well as KEGG and Panther pathways, using default settings with few modifications (sigMethod = “fdr”, fdrMethod = “BH”, fdrThr = 1, topThr = 10, reportNum = 20, perNum = 250).

#### Over Representation Analysis (ORA)

For ORA, we used the WebGestaltR package.^66^ The target list was composed of all HVGs identified for the time-point of interest, and the reference list comprised all genes expressed at that time-point. Analysis was conducted for both Biological Processes and Molecular Functions from gene ontology, as well as KEGG and Panther pathways, using default settings with few modifications (sigMethod = “fdr”, fdrMethod = “BH”, fdrThr = 0.1, topThr = 10, reportNum = 20, perNum = 1000).

#### Feature Normalization

To mitigate the influence of plate-to-plate variation, normalization prior to analysis was performed in different manners, depending on the feature type. Shape-based features were z-score normalized across the dataset. Intensity-based features were z-score normalized on a per plate basis, to mitigate plate-dependent signal differences. Lastly to visualize the radial distribution features a min-max normalization approach was taken on a per plate basis. For classification and dimension reduction z-score normalization on a per plate basis was also performed for the radial distribution features.

#### Dimensionality Reduction

Sparse PCA from scikit-learn performed on features z-score normalized across timepoints with the sparseness level alpha set to 0.7 and using 4 components. The PLSR analysis was carried out using the implementation in scikit-learn^67^, using 2 latent variables, with either developmental outcome or axis number set as target variables. Input features were z-score normalized per feature and per time-point using the normalization strategy described above. To transform the categorical annotations to a ^33^ numeric variable, successful TLSs or single axis annotations were set to 1 and unsuccessful or multiple axis annotations set to 0.

#### Classification

The state vector classifier, linear discriminant analysis, random grid search, cross-validation and scoring implementations in scikit-learn were used and the XG-Boost classifier implemented in dmlc XG boost python package was used. Any location or orientation-based features were excluded from all feature sets. Features were then z-score normalized per feature per time-point as described above. For the PLSR selected feature set used for axis classification all features within the outer 5th percentiles of loading weights of the PLSR components trained for axis number classification were selected. The correlation selected feature set used for developmental outcome classification included all features correlating with neural or somitic module scores above an absolute value of at least 0.7 and corresponding p-values below 1.0×10^−5^ were chosen for classification. To avoid test-train split bias and/or over-fitting (as can be the case for datasets, in which the feature number is close to the sample number^68^, nested cross validation was employed with stratified K-fold splits. For model performance scoring a 10-fold split was used and for hyper-parameter tuning a 3-fold split was used. Hyper-parameter tuning was performed using random grid search. Lastly, scoring was performed using the balanced accuracy score, to avoid effects due to unequal morphotype numbers (Figure 1B).

Data visualization of dimension reduction and classification results was performed using seaborn and matplotlib.^69,70^

### Code and data availability

We are in the process of depositing all sequencing data in GEO, and all imaging and related source data in Zenodo. We will update the preprint as soon as this is finished. The image-processing and phenotypic data analysis can be reproduced using the code available in https://github.com/Team-Stembryo/Integrated_Molecular-Phenotypic_Profiling_of_Stembryos and will require the functions implemented in the library https://github.com/Cryaaa/organoid_prediction_python developed for this project. Other computational code is available at https://github.com/Team-Stembryo.

## AUTHOR CONTRIBUTIONS

JVV conceived and supervised the project. RS conducted imaging data analysis, assisted by AP and AVL. AVL performed mapping of the phenotypic landscape and all functional experiments. SIG generated structures for single-cell RNA-sequencing, performed by AB. JVV performed computational analysis together with NLA and RS. ABK supervised NLA and provided critical input. SG assisted with computational analysis and functional experiments. AVL, RS and JVV wrote the manuscript, with input from all authors.

## ACKNOWLEDGEMENTS

This work was supported by the Max Planck Gesellschaft (ABK, JVV), the Sofja Kovalevskaja Award to ABK, and a Deutschen Zentrums zum Schutz von Versuchstieren (Bf3R) grant (60-0102-01.P589), EIC Pathfinder grant (Horizon-EIC-2021-PathfinderChallenges-01 101071203, SUMO), and Stiftung zur Förderung und Erforschung von Ersatz-und Ergänzungsmethoden zur Einschränkung von Tierversuchen (SET) grant to JVV. We thank Christoph Zechner, Robert Haase, Iftach Nachman, and all the members of the Veenvliet and Mateus Lab for helpful discussions. We are grateful to Stefanie Grosswendt, Christopher McGinnis and Zev Gartner for providing the MULTI-Seq reagents, and to Alysson Ryan and Carl Modes for providing early access to TosKa. We thank the Sequencing Core at the Max Planck Institute for Molecular Genetics for technical assistance, and the Organoid and Stem Cell Facility, Light Microscopy Facility, Technology Development Studio and Scientific Computing Facility at the Max Planck Institute of Cell Biology and Genetics for their outstanding continuous support.

---

**Figure S1.**
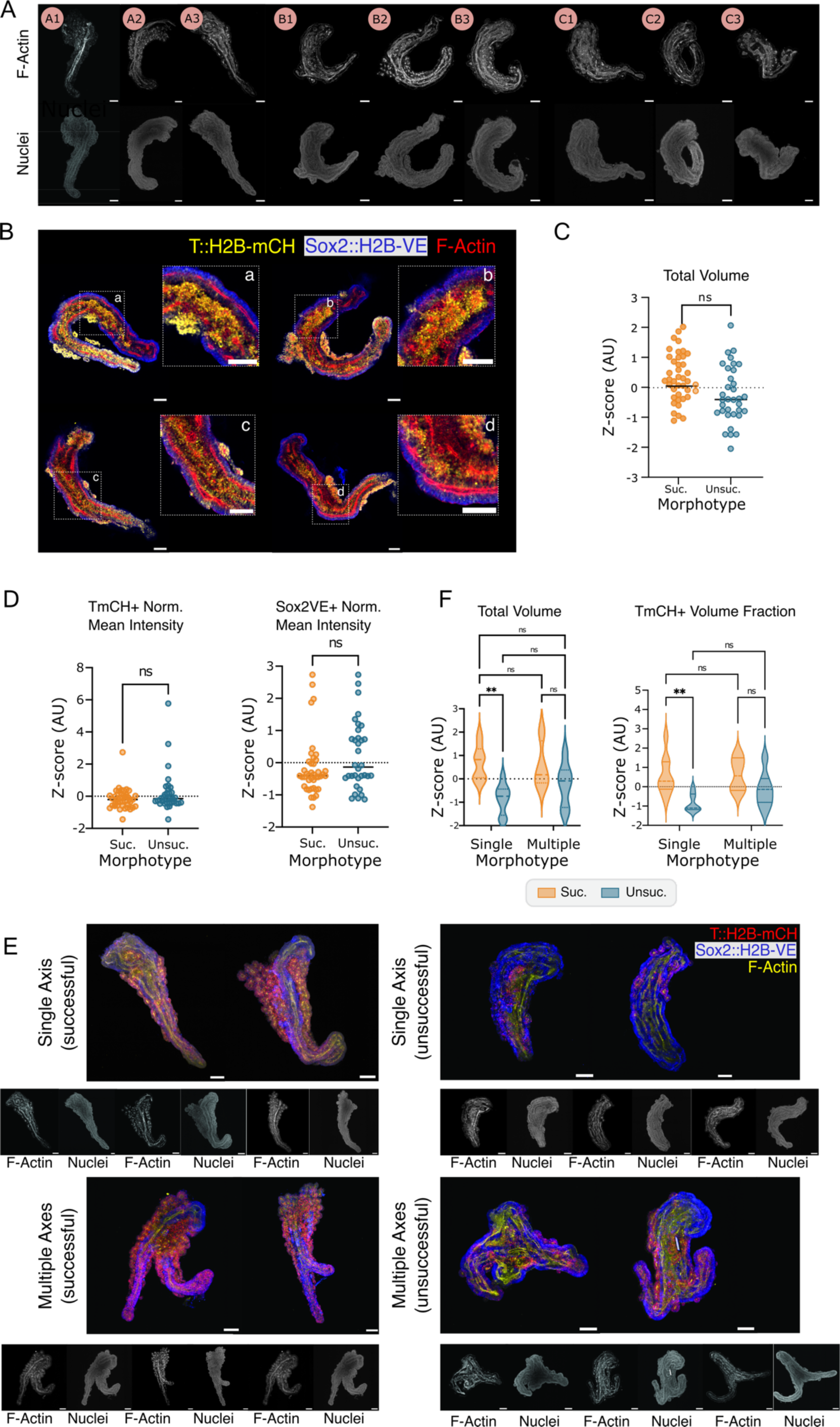
Divergent differentiation outcomes in stembryos. (A) Single planes of confocal imaging of the TLSs shown in Figure 1E, stained with Phalloidin (F-Actin, top) or DAPI (nuclei, bottom). Scale bars, 100µm (B) Composite single planes of confocal imaging of four TLSs classified as unsuccessful (based on bright-field) that display internalization of T^mCH+^ mesodermal tissue and their respective zoom-in images. The dashed lines indicate the zoom-in region. Scale bars, 100µm. (C) Total volume quantification of successful (Suc., n=41) and unsuccessful (Unsuc., n=34) unipolar TLSs. **, p < 0.01 (D) Intensity quantification of T^mCH+^ and Sox2^VE+^ signal intensity in successful (Suc., n = 41) and unsuccessful (Unsuc., n = 34) unipolar TLSs. **, p < 0.01 (E) Representative 3D maximum intensity projections (T::H2B-mCH/Sox2::H2B-VE/F-Actin composites) and single confocal planes of Phalloidin (F-Actin) or DAPI (nuclei) stained successful or unsuccessful unipolar and multipolar TLSs. Scale bars, 100µm. (F) Total volume and T^mCH+^ Volume fraction quantification of successful (Suc.) uni-(n = 9) and multipolar (n = 4) TLSs, and unsuccessful (Unsuc.) uni-(n = 8) and multipolar TLSs (n = 7). *, p<0.05, **, p<0.01, ***, p<0.001, ****, p<0.0001.

**Figure S2.**
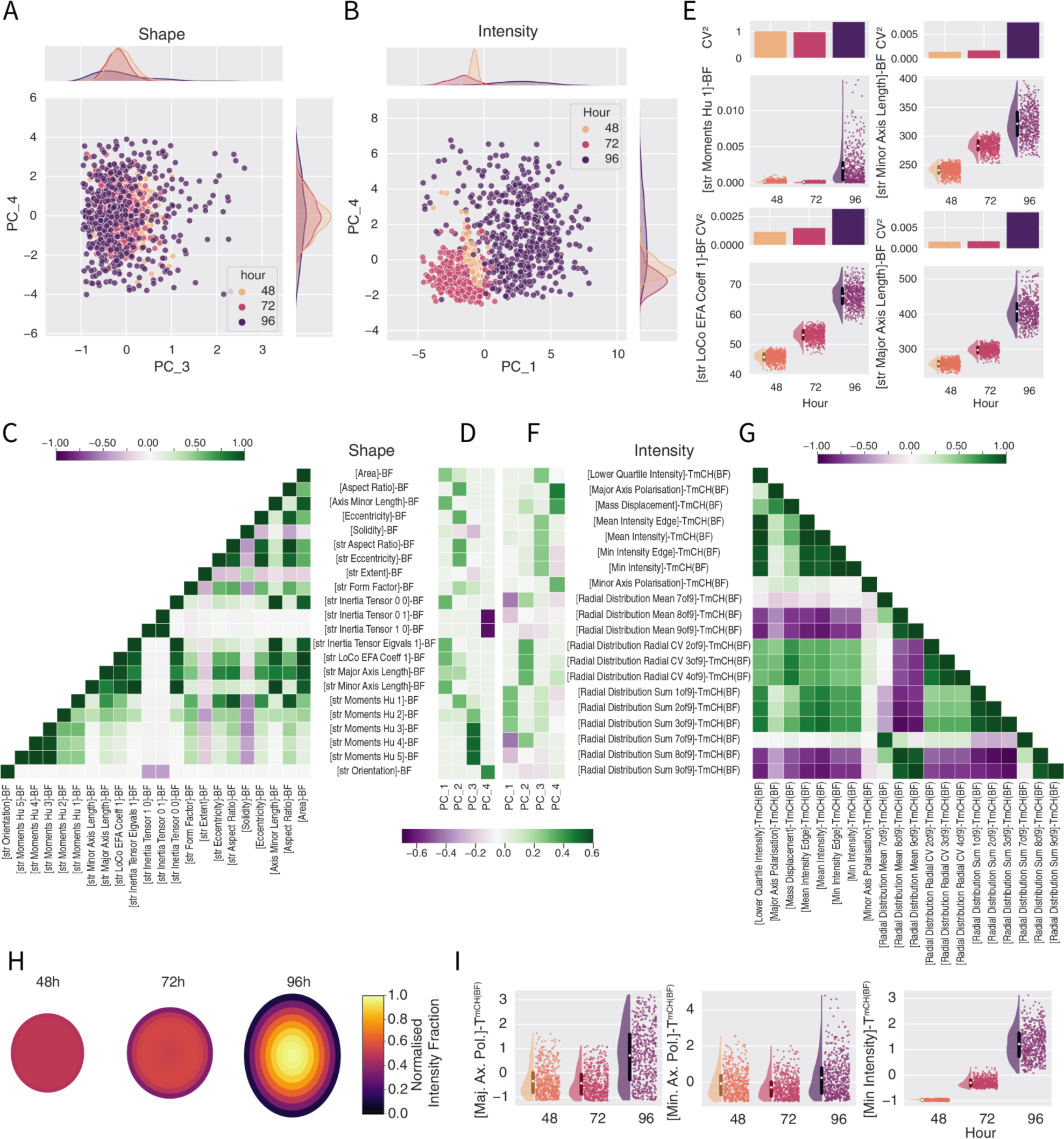
Charting of (variation in) stembryo phenodynamics. (A,B) Scatterplots of principal components determined using sparse PCA (sPCA) with BF and T^mCH^ features of all timepoints, respectively. Timepoints are highlighted to visualize the phenotypic dynamics and variation in the dataset. Top and left of the scatter-plot density plots of the x- and y-axis are shown. For a discussion of Shape PC3 and PC4 see supplemental Note 2. Sample sizes: A: 48h - n = 766; 72h - n = 662; 96h - n =546. Sample sizes B: 48h - n = 545; 72h - n = 546; 96h - n = 546. (C) Feature loadings of all principal components determined for the BF sPCA calculation. Only features that have loadings > 0.95 quantile or < 0.05 quantile are shown. (D) Correlation plot of the features shown in C. Correlation calculated using Pearson’s correlation. (E) Plots of individual features with high loadings for BF feature sPCA. Top shows CV^2^ of features at each timepoint, bottom shows combined raincloud and violin plot of feature values per timepoint. Errorbars in violin-plots show standard deviation and white center shows mean value. Sample sizes: E: 48h - n = 766; 72h - n = 662; 96h - n =546. (F) T^mCH^ features sPCA loadings. Only features that have loadings > 0.95 quantile or < 0.05 quantile are shown. (G) Correlation plot of the features shown in F. Correlation calculated using Pearson’s correlation. (I) Plots of individual features with high loadings for T^mCH^ feature sPCA. Since intensity features have to be normalized to make plates comparable, z-scores are plotted instead of raw intensities. Sample sizes: 48h - n = 545; 72h - n = 546; 96h - n = 546. (H) *[Radial Distribution Mean]*-T^mCH(BF)^ visualization. Structures are approximated as ellipses with average major and minor axes of the respective timepoints. Concentric ellipses represent the radial segments in which mean intensity fraction values are visualized by color. Sample sizes: 48h - n = 545; 72h - n = 546; 96h - n = 546. A,B,E,I: All plots visualize values where: value > 0.005 quantile and value < 0.995 quantile to reduce the influence of extreme outliers on visualization.

**Figure S3.**
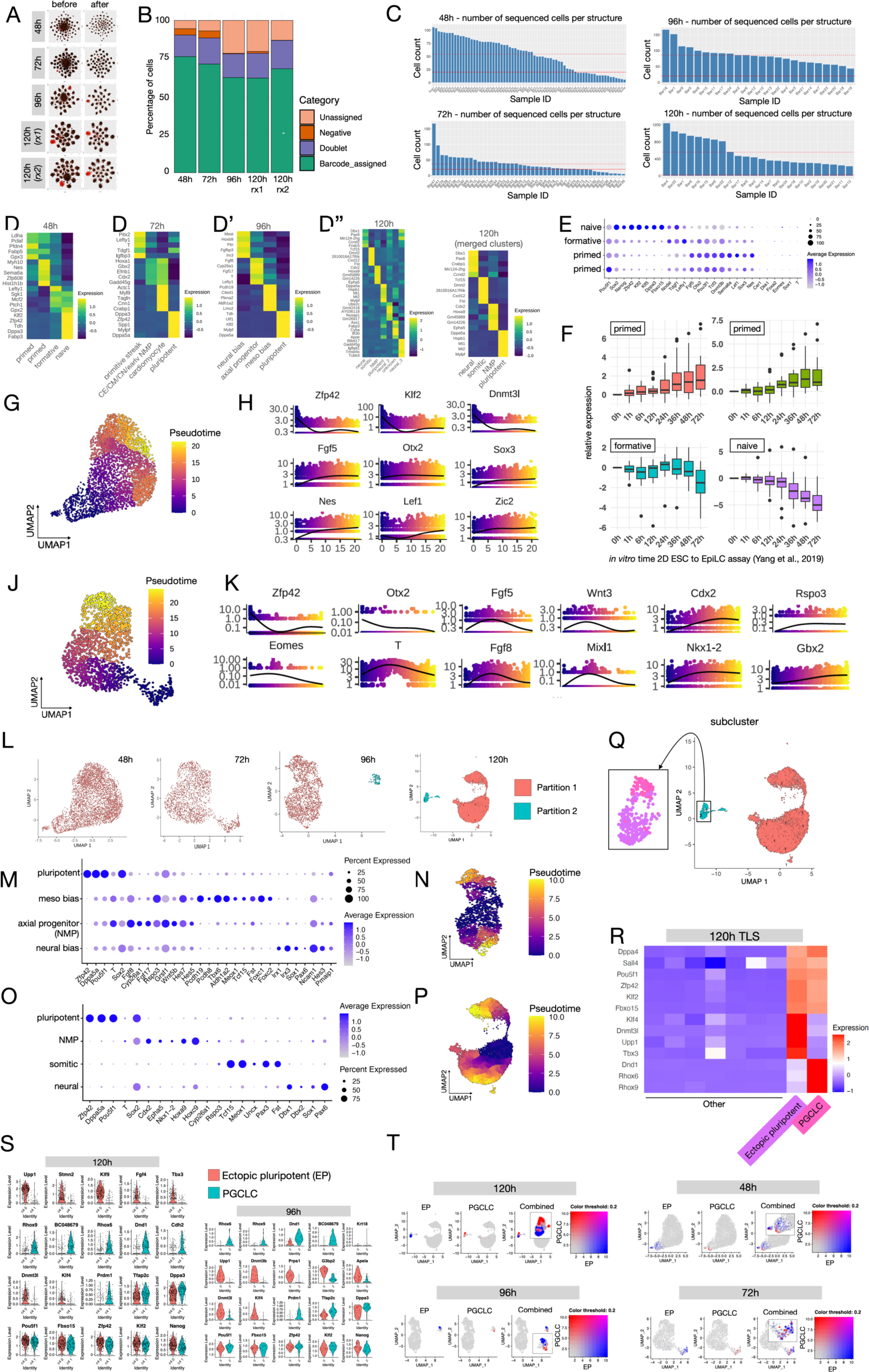
Time-resolved single-cell RNA-sequencing of individual stembryos. (A) Plots showing clustering of sample barcodes before and after removal of cells with no associated sample barcode (unassigned or negative) or more than one barcode (doublets) (see Methods). (B) Stacked barplot showing percentage of barcodes in indicated categories. Note that 120h reaction 1 and 2 (rx1/2) were integrated for downstream analyses (see Methods) [C) Histogram showing distribution of number of sequenced cells per structure at individual time-points. Dotted red line indicates the median. Solid red line represents cut-off for including sample in downstream variability analyses. (D) Heatmaps showing cluster marker genes at 48h (D), 72h (D’), 96h (D’’) and 120h (D’’’). For 120h, an additional heatmap is shown, where the three neural clusters were pooled (D’’’ right). (E) Dot Plot showing expression of established naïve, formative, and primed pluripotency genes, as well as lineage factors. (F) Boxplot showing the distribution of the expression of cluster marker genes at the indicated time-points in the *in vitro* 2D ESC to EpiLC differentiation assay (see Methods).^25^ (G) 48h UMAP with cells colored by pseudo-time. (H) Expression of selected genes as a function of pseudo-time. (I) Dot Plot showing expression of established pluripotency, caudal epiblast (CE) / caudal mesoderm (CM), caudal neuro-ectoderm (CN) and primitive streak marker genes. (J) 72h UMAP with cells colored by pseudo-time. (K) Expression of selected genes as a function of pseudo-time. (L) UMAPs of individual time-points, colored by partitions. (M) Dot Plot showing expression of pluripotency, mesodermal, neural and axial progenitor (NMP, neuro-mesodermal progenitor) genes. (N) 96h UMAP with cells colored by pseudo-time (only partition 1). (O) Dot Plot showing expression of pluripotency, somitic, neural and NMP genes. (P) 120h UMAP with cells colored by pseudo-time (only partition 1). (Q) Sub-clustering of 120h, partition 2 (cyan in 120h UMAP) reveals two distinct cell populations (insert) (R) Heatmaps showing distinct transcriptomic signatures of two sub-clusters: ectopic pluripotent cells (EPCs) and primordial germ cell like cells (PGCLCs) (S) Violin Plots showing expression of *in vivo* naïve pluripotency and PGC markers^25,71^ in EPCs and PGCLCs. (T) UMAPs of individual time-points, colored by EPC module score (blue), PGCLC module score (red) or combined. Inserts show clear separation of both populations over time (for calculation of module scores, see Methods).

**Figure S4.**
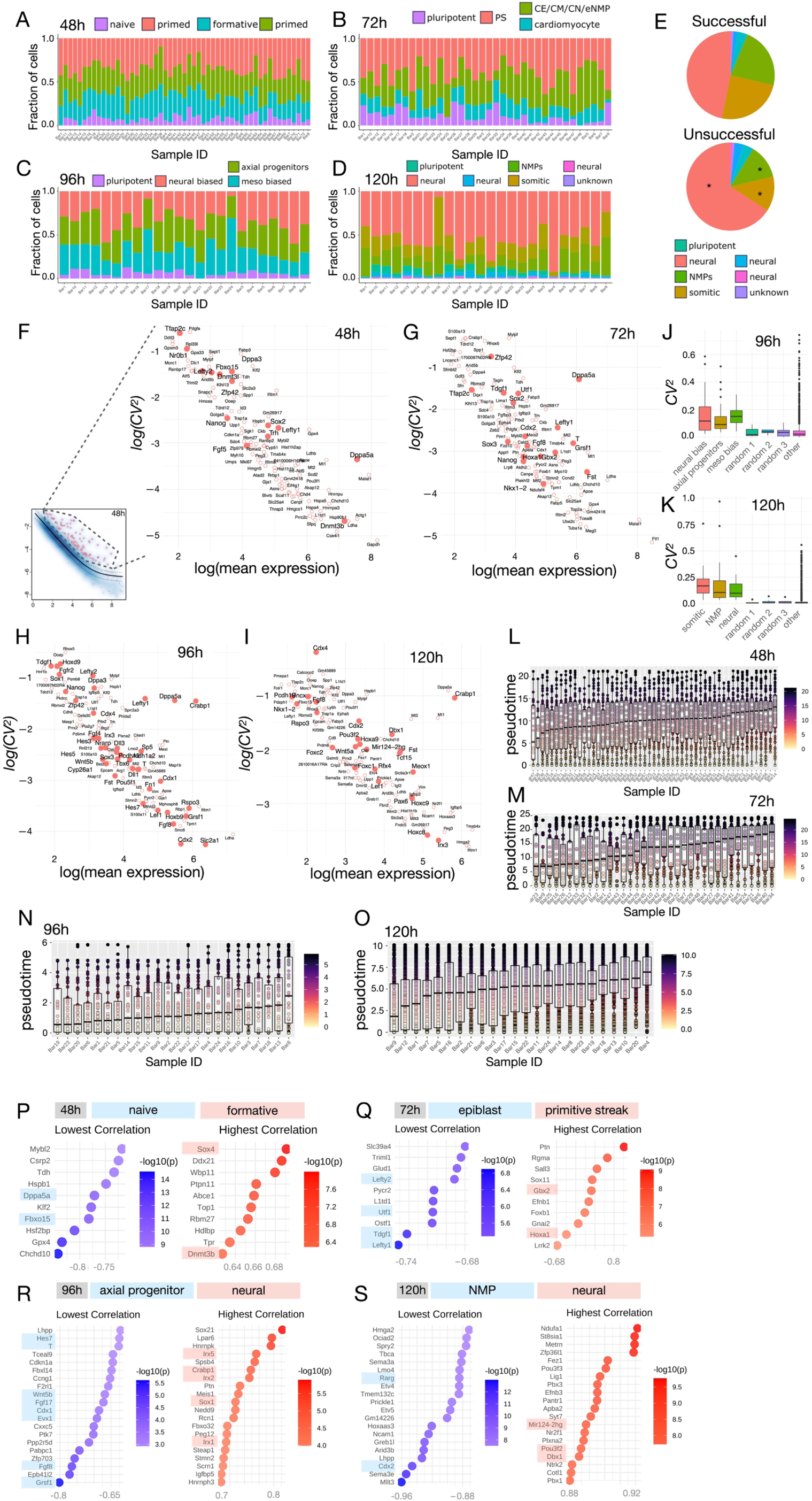
Profiling of inter-stembryo variation in transcriptional signatures. (A-D) Stacked bar plots showing cellular composition of individual structures at indicated time-points. (E) Pie charts showing distribution of cell states in successful and unsuccessful TLSs. *, p < 0.05 (ANOVA with BH FDR correction). (F) Plot showing top 100 HVGs at 48h (red dots in full plot bottom left). (G-I) Plot showing top 100 HVGs at 72h (G), 96h (H), 120h (I). (J,K) Boxplots showing distribution of *CV*^2^ values for marker genes of indicated clusters, randomly selected or all other expressed genes at 96h (J) and 120h (K). (L-O) Boxplots showing distribution of pseudotime values in individual structures at indicated time-points. (P-S) Plots showing top 10 (48h (P) & 72h (Q)) or top 20 (96h (R) & 120h (S)) genes that have the lowest (blue) and highest (red) correlation with average pseudotime.

**Figure S5.**
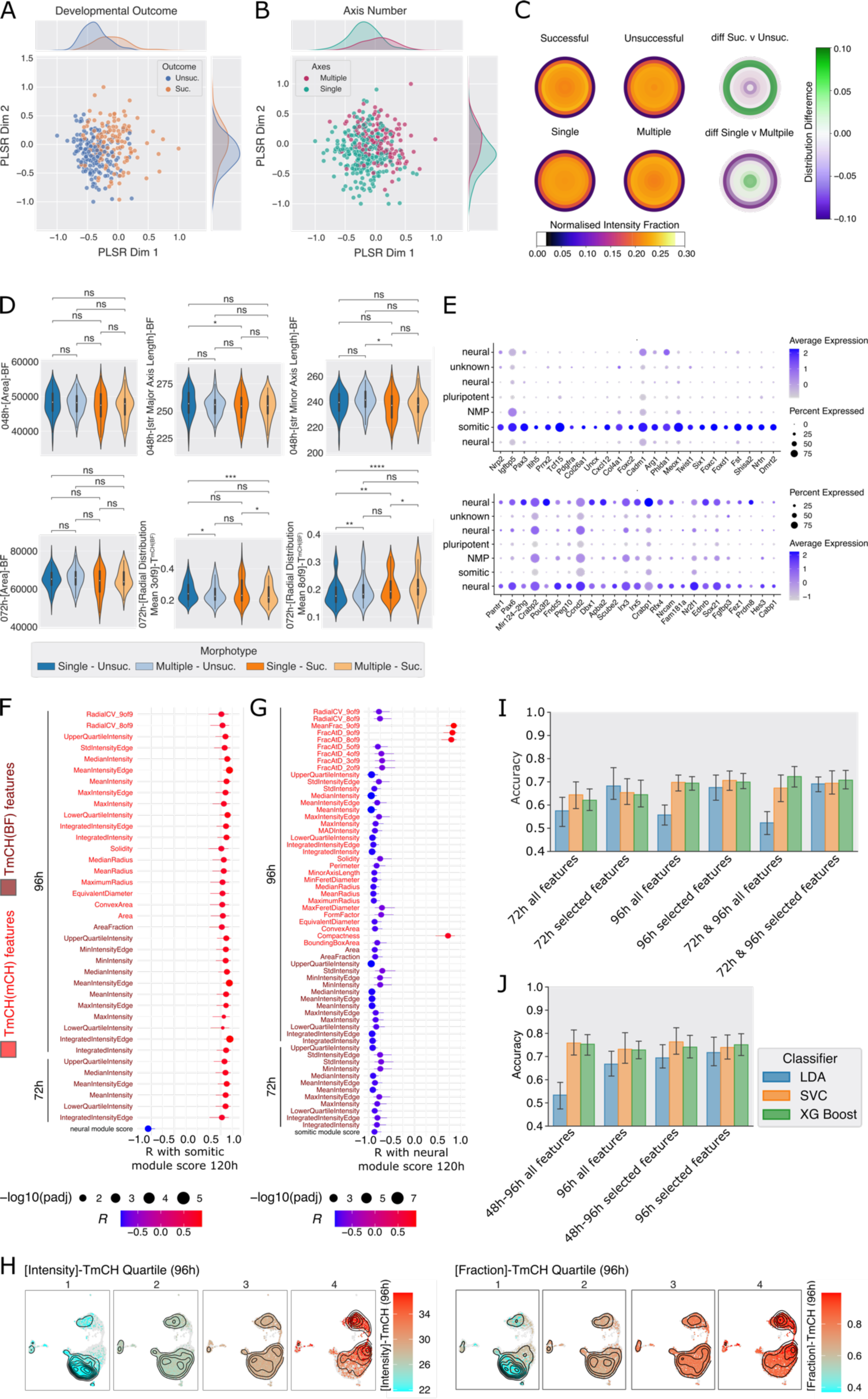
Identification and validation of features predictive of stembryo differentiation outcome. (A,B) Scatterplots of PLSR latent variables trained with developmental outcome and axis number as target variables, respectively, using only features of structures at 48h and 96h. To the top and right of the plots the density plots of the x and y axis values, respectively are shown. Successful TLS development: Suc; unsuccessful TLS development: Unsuc. Sample sizes A: Unsuccessful – n = 183, Successful – n = 128. Sample sizes B: Multiple – n =136; Single – n = 256. (C) : 072h-*[Radial Distribution of CV]*-T^mCH(BF)^ visualization. Ellipses approximate the shape of structures with average major and minor axes of the respective morphotypes. Inner concentric ellipses represent the radial segments in which the normalized intensity fraction is visualized by color. Distribution difference visualization: inner concentric ellipses represent the radial segments in which the difference between morphotypes is visualized by diverging colors. Sample sizes: Unsuccessful – n = 183, Successful – n = 129, Multiple - n =136; Single – n = 257 (D) : Individual feature plots, plotted for all combinations of morphotypes. * p < 0.05, ** p < 0.01, *** p < 0.001, **** p < 0.0001. *[feature]*-T^mCH(BF)^: z-scores of values plotted to take into account plate to plate variation (see methods). Sample sizes: Multiple-Unsuccessful – n = 55; Multiple-Successful – n = 34; Single-Unsuccessful – n = 128; Single- Successful – n =95. (E) Dot Plot showing expression of top 25 somitic (top) and neural (bottom) markers in all 120h clusters. These markers were used to calculate somite and neural module scores respectively (F,G) Plot showing all features significantly (Padj < 0.05) correlated with 120h somite (F) or neural (G) module score. Dot indicates correlation with bars representing Confidence Interval. Dot size scales with -log10(Padj). (H) Dim Plots showing distribution of 120h sequenced cells per 96h [Intensity]-T^mCH^ quartile (left) or [Fraction]-T^mCH^ quartile (right). All cells are shown in gray. Contours show relative frequency of data. (I) Classifier balanced accuracy scores shown developmental outcome prediction using several feature sets. All Features: complete feature set; Corr. Sel. Features: features significantly correlating with somitic or neural module scores. 72h, 96h, 72h & 96h indicates from which timepoint features are included in the set. (J) Classifier balanced accuracy scores for axis number prediction using several feature sets. All Features All Timepoints: complete feature set; PLSR Selected Features: all features that were outside of the 90th percentile of loading weights, the corresponding 96h feature sets are identical but all non-96h features are removed (see methods for details on all feature sets).

**Figure S6.**
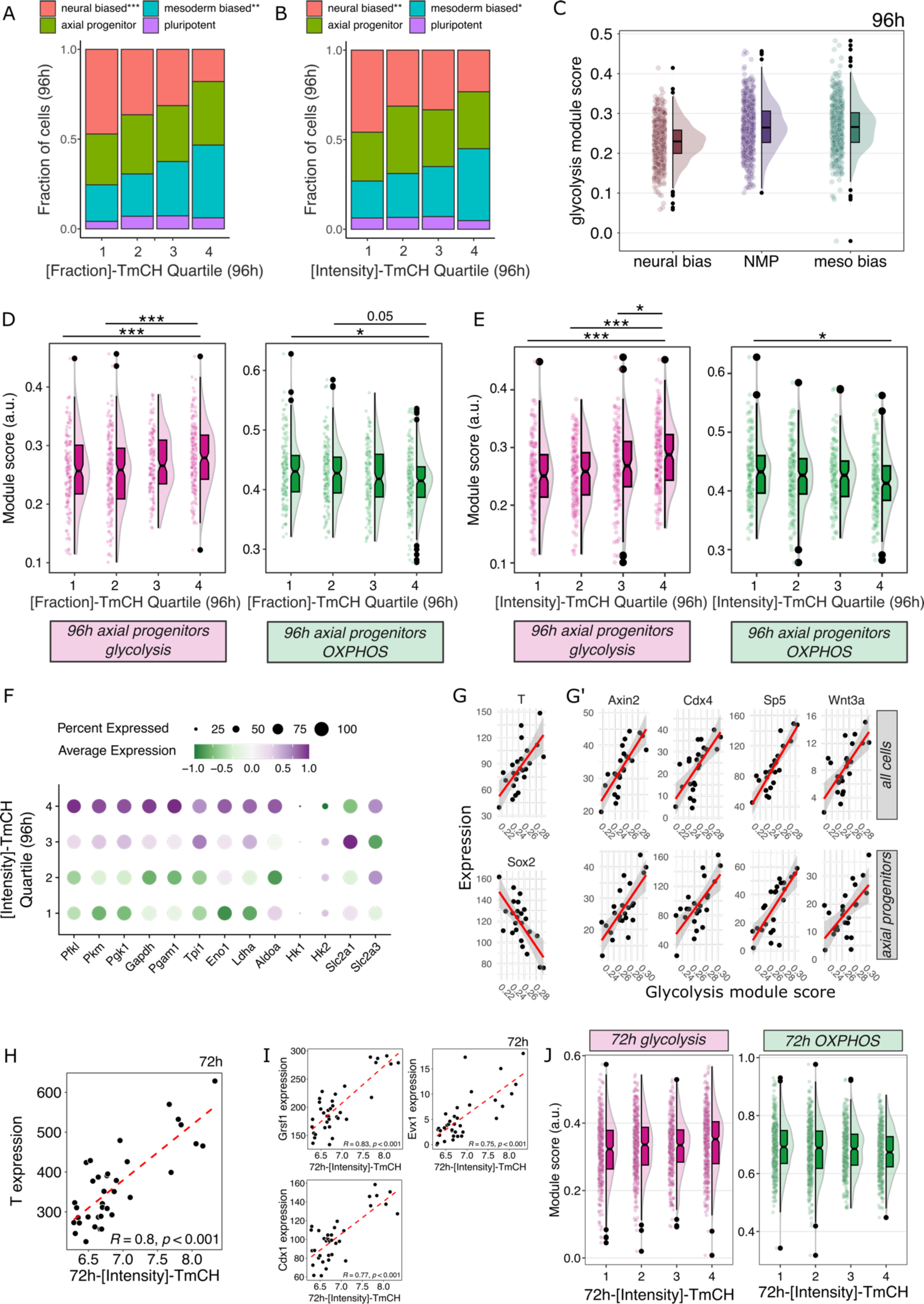
Identification of biological processes underlying phenotypic variation. (A,B) Cumulative bar plot showing the fraction of cells in each state for [Fraction]-T^mCH^ quartiles (A) and [Fraction]-T^mCH^ quartiles (B) at 96h. * p<0.05, ** p<0.01, *** p<0.001 (ANOVA with BH FDR correction). (C) Raincloud plots showing distribution of glycolysis module score in indicated cell clusters. (D,E) Raincloud plots showing glycolysis module score (magenta) and OxPhos module score (green) per [Fraction]-T^mCH^ quartile (D) or [Intensity]-T^mCH^ quartile (E) at 96h. * padj<0.05, ** padj<0.01, *** padj<0.001. (F) Dot plot showing average expression of indicated genes encoding glycolytic pathway genes per [Intensity]-T^mCH^ quartile at 96h in NMPs. (G,G’) Scatter plot showing correlation of expression of indicated genes and glycolysis module score at 96h in all cells (top) or NMPs (bottom). Black dots are individual samples. Grey area indicates 95% CI. (H) Scatter plot showing correlation of *T* expression and [Intensity]-T^mCH^ quartile at 72h. Black dots are individual samples. (I) Scatter plot showing correlation of *T* expression and [Intensity]-T^mCH^ quartile at 72h. Black dots represent individual samples. (J) Raincloud plots showing glycolysis module score (magenta) and OxPhos module score (green) per [Intensity]-T^mCH^ quartile at 72h.

**Figure S7.**
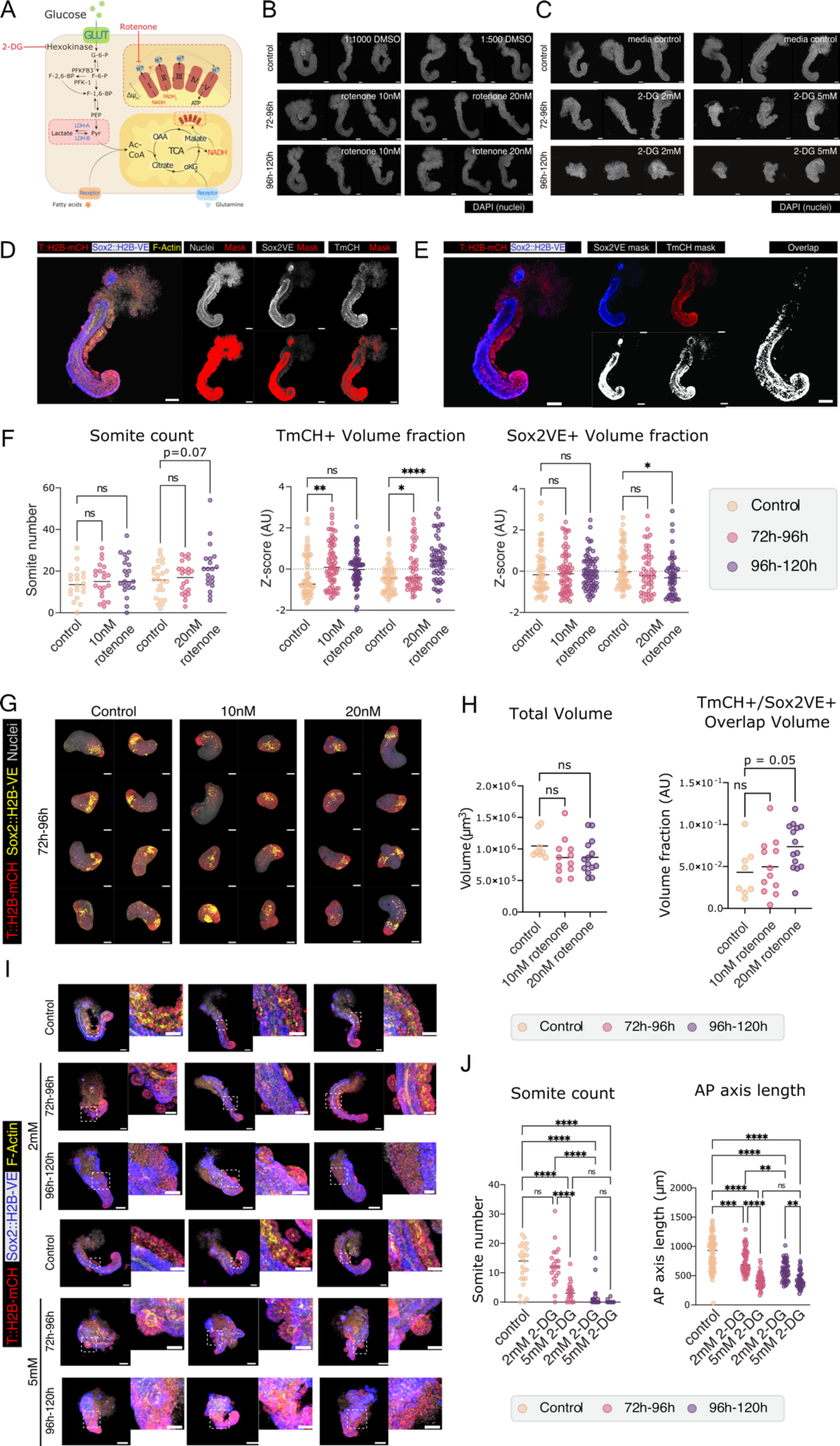
Identification and modulation of biological processes associated with divergent differentiation outcomes. (A) Schematic representation of glucose metabolism pathways. The metabolic flux of glucose can be modulated using different inhibitors. Inhibitors in red designed to alter OXPHOS-glycolysis balance (see text). (B,C) Representative 3D maximum intensity projections (3D-MIPs) of DAPI images obtained by confocal imaging of 120h TLSs treated with rotenone (B) or 2-DG (C) during indicated time intervals, and their respective controls. Scale bars, 100 µm. (D,E) Example of threshold used to establish the masks to quantify T^mCH+^ and Sox2^VE+^ intensity (D) and (overlap) volumes (E) (See Methods). Scale bars, 100 µm. (G) Quantifications of Somite Count (n=20 TLSs per condition), T^mCH+^ and Sox2^VE+^ Volume Fraction of 120h TLSs treated with Rotenone 10nM (72h-96h, n=69; 96h-120h, n=53) or 20nM (72h-96h, n=46; 96-120h, n=55), and their respective controls (Control for 10nM, n=59, Control for 20nM, n=63). *, p<0.05, **, p<0.01, ***, p<0.001, ****, p<0.0001. (H) Representative 3D-MIPs of confocal imaging of 96h gastruloids treated with 10nM and 20nM Rotenone and their respective controls. Scale bars, 100 µm. (I) Quantification of the Total Volume and T^mCH+^ and Sox2^VE+^ Overlap Volume Fraction of 96h structures treated with 10nM (n= 12) and 20nM (n=14) Rotenone from 72-96h, and their respective control (n=8). *, p<0.05, **, p<0.01, ***, p<0.001, ****, p<0.0001. (J) Representative 3D-MIPs of 120h TLSs treated with 2-DG during indicated time intervals, and controls. Area indicated by white dashed is shown in close-ups to highlight somite morphology and architecture under different 2-DG treatments. Scale bars, 100µm. (K) Quantifications of Somite count (n=20 per condition) and Major (AP, Anterior-Posterior) axis length of 120h TLSs treated with 2-DG 2mM (72h-96h, n=100; 96h-120h, n=105) or 5mM (72h-96h, n=100; 96-120h, n=965), compared to control (n=220). *, p<0.05, **, p<0.01, ***, p<0.001, ****, p<0.0001.

## SUPPLEMENTARY TABLES

**Supplementary Table 1. PLSR Loadings and Feature Correlation Matrix**

**PLSR Loadings (All Timepoints):** Feature loadings of PLSR latent variables (Fig. 3A, B). Features shown are in the top 95th percentile or bottom 5th percentile of loadings for either the developmental outcome PLSR latent variables or axis number PLSR latent variables. Loadings colored by a diverging colormap to aid in interpretability. **PLSR Loadings (Early Timepoints):** Feature loadings of PLSR latent variables for PLSR analysis performed using only early features (48h and 72h - Fig. S5A, B). Features shown are in the top 95th percentile or bottom 5th percentile of loadings for either the developmental outcome PLSR latent variables or axis number PLSR latent variables using only early features. Loadings colored by a diverging colormap to aid in interpretability. **Correlation Matrix:** Correlation Matrix of all features shown in Tab: PLSR Loadings (All Timepoints). Correlation displayed is Pearssons correlation. All correlation values colored by a diverging colormap to aid in interpretability.

**Supplementary Table 2.**
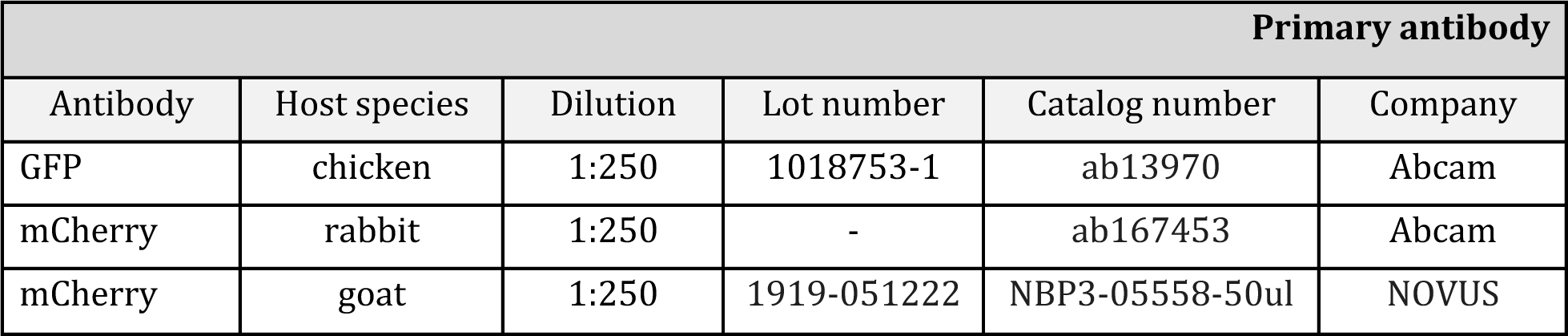
List of Antibodies used in this study.

| Primary antibody |  |  |  |  |  |
| --- | --- | --- | --- | --- | --- |
| Antibody | Host species | Dilution | Lot number | Catalog number | Company |
| GFP | chicken | 1:250 | 1018753-1 | ab13970 | Abcam |
| mCherry | rabbit | 1:250 | - | ab167453 | Abcam |
| mCherry | goat | 1:250 | 1919-051222 | NBP3-05558-50ul | NOVUS |

**Supplementary Table 3.** List of MULTI-seq oligonucleotide sequences and corresponding sample IDs.

| Secondary antibody/Conjugated dyes |  |  |  |  |  |
| --- | --- | --- | --- | --- | --- |
| Antibody | Wavelength | Dilution | Lot number | Catalog number | Company |
| DAPI | 405 | 1:20000 | 28718-90-3 | 10236276001 | Merck Roche |
| Donkey-anti-chicken | Alexa 488 | 1:250 | 162189 | 703-545-155 | BIOZOL |
| Donkey-anti-goat | Alexa 546 | 1:500 | A11056 | 2306813 | TFS |
| Donkey-anti-rabbit | Alexa 568 | 1:500 | 2540901 | A10042 | TFS |
| Phalloidin | 647 | 1:250 | 2431325 | A22287 | Invitrogen |
| Phalloidin | 647+ | 1:250 | 2585754 | A30107 | TFS |

## SUPPLEMENTAL NOTE 1

### Feature descriptions

To gain some intuition into what the features used during the analysis of wide-field microscopy describe, we have compiled some short descriptions (**Table SN1,2**) and selected visualizations (**Figure SN1**). In Tab. 1 short descriptions of shape-based features are shown, whereas Tab. 2 contains descriptions of intensity-based features.

**Table SN1.**
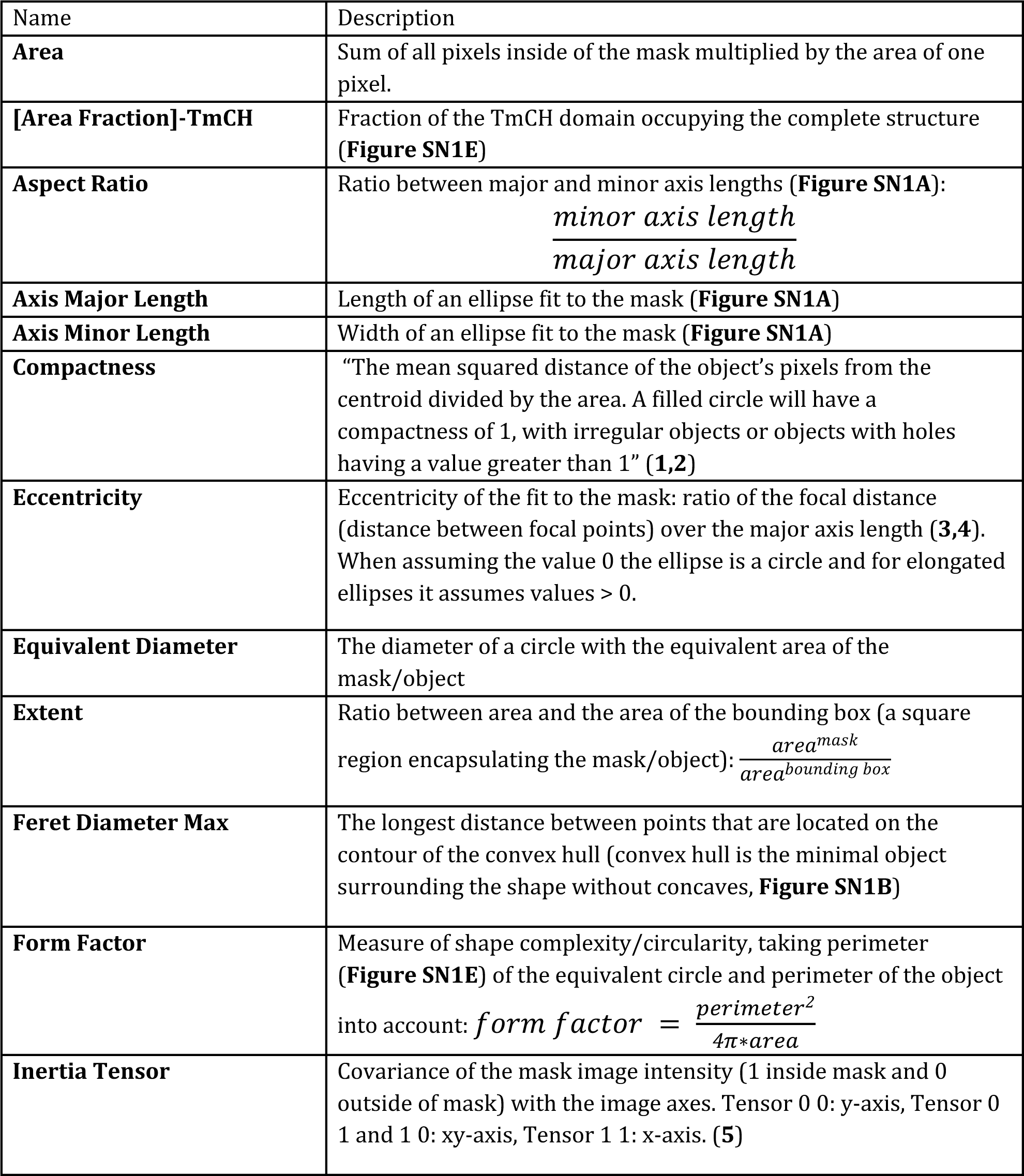
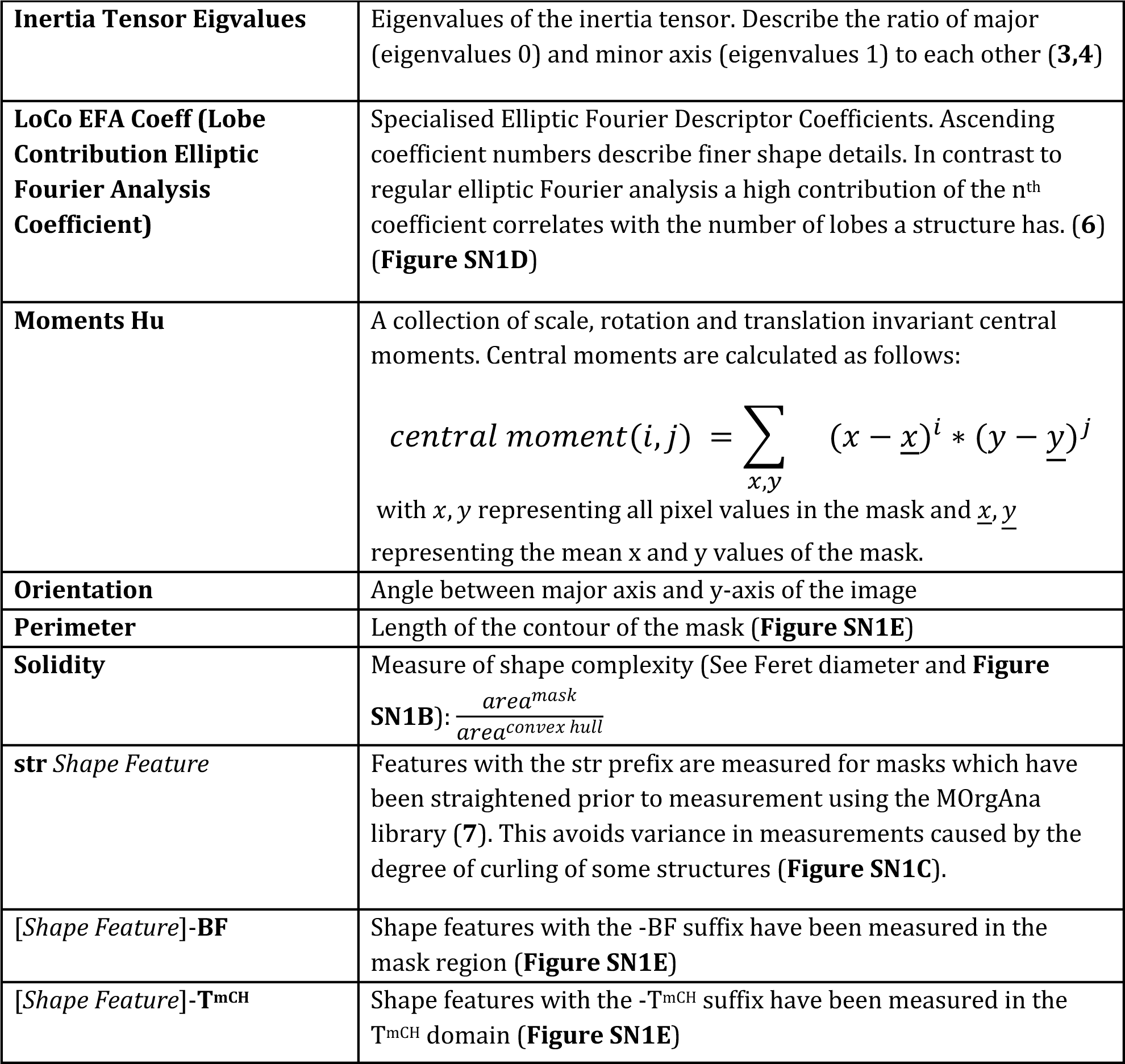
Shape Feature Description: Short descriptions of shape features.

**Figure SN1.**
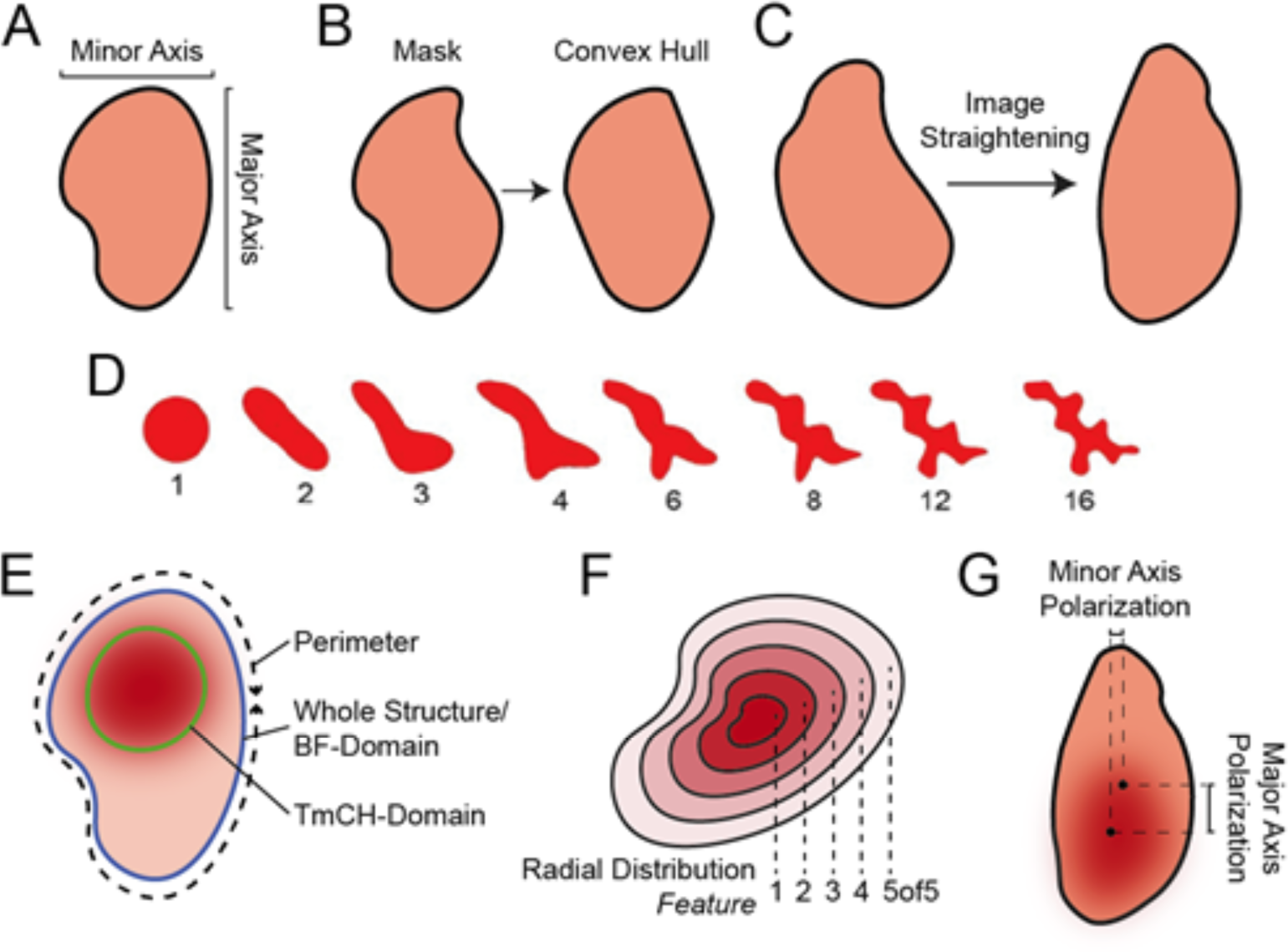
Visualization of Selected Features: **A:** Visualization of major and minor axis lengths. Aspect ratio is the ratio of both. **B:** Convex hull of a mask. **C:** Image straightening implemented in MOrgAna shown for an example mask. **D:** Shape of an object reconstructed using the first n LOCO-EFA coefficients. Modified from (7) under a Creative Commons 3.0 license. **E:** Visualization of BF and T^mCH^ domains, as well as the perimeter of the BF-domain. **F:** Visualization of the radial distribution features. **G:** Visualization of major and minor axis polarization metric.

**Table SN2.** Intensity Feature Description: Short descriptions of intensity features.

| Name | Description |
| --- | --- |
| **Integrated Intensity** | Sum over all intensity values of all pixels within the mask/domain |
| **Lower Quartile Intensity** | Intensity value of the 25<sup>th</sup> percentile |
| **MAD Intensity** | Median Absolute Deviation calculated inside of the mask/domain, i.e. the average difference between all pixels and the Median Intensity |
| **Mass Displacement** | Distance between mask/domain centroid and weighted centroid (weighted by intensity value) inside of the mask/domain (1,2) |
| **Max Intensity** | Maximum intensity value of all pixels within the mask/domain |
| **Mean Intensity** | Mean intensity value of all pixels within the mask/domain |
| **Median Intensity** | Median intensity value of all pixels within the mask/domain |
| **Min Intensity** | Minimum intensity value of all pixels within the mask/domain |
| **Std Intensity** | Standard Deviation of all pixel intensity values within the mask/domain |
| **Upper Quartile Intensity** | Intensity of the 75<sup>th</sup> percentile |
| **Radial Distribution Sum Nof9** | Fraction of the summed intensity in the region Nof9 (**Figure SN1F**) to the integrated intensity of the whole mask/domain. |
| **Radial Distribution Mean Nof9** | Radial Distribution Sum normalised by the area of the region Nof9 (**Figure SN1F**) |
| **Radial Distribution Radial CV Nof9** | Coefficient of variation calculated inside region Nof9 (**Figure SN1F**) |
| **Major Axis Polarisation** | Distance along the major axis between the centroid of the mask and the weighted centroid of the region of highest intensity (values ≥ 95<sup>th</sup> percentile intensity) (**Figure SN1G**) |
| **Minor Axis Polarisation** | Distance along the minor axis between the centroid of the mask and the weighted centroid of the region of highest intensity (**Figure SN1G**) |
| *Intensity Feature Edge* | *Intensity Feature* measured for pixels belonging to the contour of the mask/object |
| *[Intensity Feature]-T<sub>mCH</sub>(BF)* | *Intensity Feature* measured on all pixels within the structure boundary (**Figure SN1E**) |
| *[Intensity Feature]-T<sub>mCH</sub>(mCH)* | *Intensity Feature* measured on all pixels within the T<sub>mCH</sub> domain boundary (**Figure SN1E**) |

## SUPPLEMENTAL NOTE 2

In our SPCA analysis using shape-based, PC3 and PC4 do not separate timepoints, yet the 96h structures seem to have a high variance for these components (**Figure S2A)**. The loadings for these components are dominated by inertia tensors, hu moments, solidity, extent and orientation (**Figure S2D**). Especially in PC4 we observed almost exclusively contributions from inertia tensors and orientation. Since inertia tensors can be rotation invariant and share high loadings with orientation, this suggests that this component is capturing arbitrary variation, possibly due to distinct positioning under the microscope. PC3 on the other hand has high contributions of Hu moments, extent and solidity, variation in which increases upon the emergence of more complex shapes (**Figure S2C,E**).

In our SPCA analysis using T^mCH^ features, PC1 separated 72h and 96h samples from 48h samples assuming values between the two timepoints with very little variation inside the 48h group (**Figure S2B**). The loading for this component is clearly governed by radial distribution features (Radial Distribution Mean and Sum, which highly correlate with each other (**Figure S2F,G**)). When inspecting *[Radial Distribution Mean 9of9]*-T^mCH(BF)^ more closely (**Figure S2H**) we observed that an initially mostly uniform distribution of intensity at early time-points becomes more polarized towards the over time. It is possible that the high loading weight of *[Radial Distribution Mean 7of9]*-T^mCH(BF)^ causes the peculiar separation of timepoints as for this region values in 48h and 96h samples are more similar than in 72h and 96h samples (**Figure S2H**).

